# An effective model for community-based conservation around authorized fishing settlements inside a devolved Wildlife Management Area in southern Tanzania

**DOI:** 10.1101/2024.07.30.605829

**Authors:** Lily M Duggan, Lucia J Tarimo, Katrina A Walsh, Deogratius Roman Kavishe, Ramiro D Crego, Manase Elisa, Felister Mombo, Fidelma Butler, Gerry Killeen

## Abstract

Wildlife Management Areas (WMAs) represent a relatively new institutional model for devolved, locally-led conservation in Tanzania, in which local villages set aside part of their land for wildlife conservation and manage that resource collectively, so that their stakeholder communities can collectively leverage economic and social benefits from income-generating activities like tourism. This study examines the relationship between community-defined land use plans and *de facto* land use practices, and the influence of the latter on the relative abundance and distribution of large wild mammals in a across the Ifakara-Lupiro-Mangula (ILUMA) WMA, which acts as a key buffer zone between Nyerere National Park (NNP) to the east and adjacent stakeholder villages to the north and west. All observed signs of wildlife and human activity were recorded across 32 locations inside ILUMA and in the permanent settlements and national park that respectively border it to the west and east. Across much of ILUMA WMA, in areas where agreed land use plans were not adhered to, rampant cattle herding and land clearing for agriculture were associated with reductions in wildlife richness and biodiversity, as well as overall ecosystem integrity. Although human settlement was also generally associated with reduced natural ecosystem integrity, some important exceptions to this rule illustrate how sustainable livelihoods for local people that are based on well-managed natural resource harvesting practices may actually enhance conservation effectiveness: Three authorised human settlements within the WMA, where fishing was the primary permitted livelihood and local communities collaborated with the WMA management, were surrounded by pristine land cover with thriving terrestrial wildlife populations. Correspondingly, the best conserved parts of the WMA not only included those closest to the boundary with the national park to the east, but also these fishing villages along the riverbank to the north, where compliance with agreed land use plans was most rigorous. Overall, this study documents a useful example of how a devolved conservation area may conditionally host resident local communities undertaking selective natural resource extraction activities and collaborate with them to achieve effective *de facto* conservation practices.

## 1 INTRODUCTION

Locally devolved community-based conservation (CBC) is considered a promising alternative to traditional fortress-style conservation, many different forms of which have been developed in dozens of countries around the world, with varying degrees of success and/or failure. Two outcomes that are considered successes for CBC are the conservation of natural ecosystems and the many species that live in them, and participation by the local communities, not only in the governance and management of the area, but also in the derived economic and social benefits (Songorwa, 1999). By providing tangible economic benefits to local communities, through a participatory approach to land use planning and management (Gibson and Marks, 1995). The overall idea of CBC is that protected landscapes and natural resources are managed in a devolved manner by empowered local communities, to secure their future on a sustainable basis (Gibson and Marks, 1995). The aim of CBC is to shift the focus of conservation from exclusively centralized, state-controlled protection models, like national parks and game reserves, to management through bottom-up institutional structures that ensure community participation and inclusion (Goldman, 2003). In principle, local communities can benefit from CBC via consumptive tourism, such as regulated trophy hunting (Lindsey et al., 2006) and non-consumptive eco-tourism (Kiss, 2004), both of which aim at strengthening conservation efforts by keeping financial rewards within the local economy, including sustainable livelihoods for community members.

Having said that, many CBC initiatives have failed achieve these objectives, and this has often been attributed to a lack of business, tourism and marketing skills among the local stakeholder community (Spenceley, 2012). Furthermore, efforts to develop CBC practices across Africa during the 1990s have been criticised for sidelining local communities from the processes of developing management plans, while nevertheless including the community in implementing externally-imposed conservation policies (Agrawal and Ribot, 1999, Goldman, 2003). In essence, while relevant communities were, in theory, included in wildlife conservation policies, meaningful devolution of power was not implemented in practice, leading to a lack of governance stakeholdership and management authority at a local level. This prevented relevant communities from playing an active role in conservation or accrue the derived socio-economic benefits that are so important to sustain enthusiasm for such initiatives (Agrawal and Ribot, 1999).

It is therefore unsurprising that various CBC initiatives across Africa have yielded a mixed picture of success and failure (Salerno *et al*., 2016). With pressure on land use only expected to increase in the coming decades, with a predicted 9.7 billion global citizens by 2050 (Population Division of the United Nations Department of Economic and Social Affairs, 2022). Africa in particular is expected to see declines in wild mammal habitat of up to 25% by 2050, the primary driver of this being land-use change (Baisero et al., 2020). There is therefore an urgent need to establish more effective mechanisms for sustained CBC that benefit both people and wildlife as scalable models of best practice.

The establishment of Wildlife Management Areas (WMAs), as an institutional mechanism for CBC in Tanzania, is correspondingly intended to satisfactorily devolve power and rectify these management and governance issues. Although mentioned in the Wildlife Policy of 1998, WMAs were only established formally through the *Establishment and Management of Wildlife Management Areas and Benefit Sharing* Government Notice in The Wildlife Conservation Act of 2005, which was then revised in 2022 (United Republic of Tanzania Ministry for Natural Resources and Tourism, 2022). It is the intention of this institutional basis for CBC is to “empower local communities to be more involved and to hold more authority over the management of wildlife to promote sustainable biodiversity conservation and rural economic development” (USAID, 2018). In accordance with this act, WMAs should establish administrative mechanisms whereby wildlife conservation is promoted, economic development of the stakeholder villages is enhanced, rural poverty is reduced, and the distribution of benefits and costs is equitable (United Republic of Tanzania Ministry for Natural Resources and Tourism, 2022).

Each WMA has its own individual set of by-laws and is independently managed. Once formally established and authorized, the community-based management body of the WMA has the right to negotiate and sign agreements with customers and investors, under the condition that representatives of the district council are present and involved in the negotiation, with the aim of advancing both community development and wildlife conservation objectives. The WMA is monitored for unauthorised human activities by Village Game Scouts (VGS), who are appointed by the WMA management as its equivalent of wildlife rangers. VGS are also responsible for both protecting the natural resources of the area and also protecting people’s lives and property from wildlife. The VGS and WMA management also collaborate with other legal bodies, such as the Tanzanian National Parks Authority (TANAPA), The Wildlife Division, the police and the district government in operations to prevent various forms of encroachment upon protected land. Currently 19 WMAs are operational in Tanzania, accounting for 7% of the country’s total land area, and a further 19 were in the planning stages as of 2018 (Lee and Bond, 2018).

In May 2015, the Ifakara-Lupiro-Mangula (ILUMA) WMA was issued user rights and declared the 19^th^ WMA in Tanzania (United Republic of Tanzania Ministry of Natural Resources and Tourism, 2015). Fourteen villages from both the Ulanga and Kilombero districts, with a total of more than 80,000 residents, are included as stakeholder communities in this WMA (United Republic of Tanzania Ministry of Natural Resources and Tourism, 2015). Three authorised local cooperative fishing communities are established and located inside the wetland zone of ILUMA; *Mikeregembe*, *Mdalangwila* and *Funga*. In the management plan these are referred to as *Beach Management Units* (BMUs). Monitored fishing is permitted in these areas under licence and the ILUMA by-laws include regulations on how these settlements are maintained. For example, construction of a permanent residence in these locations is not permitted, so all structures must be temporary in nature. Specifically, this means that they are built using only natural materials like mud, sticks and thatch, rather than bricks and iron sheets. The residents of these areas are allowed to collect dry wood for such purposes as cooking and drying fish but are not permitted to harvest any live trees. Charcoal burning is also not permitted.

At face value, these guidelines and management mechanisms appear ideal for protecting both the livelihoods of the local people from the 14 villages and conserving wildlife. However, anecdotal evidence and informal experiences of some of the authors circa 2018 suggested that the WMA was struggling, and that the land use plans formally agreed upon were not being adhered to. The aim of this study was therefore to evaluate how well the formally agreed land use plans compared with the *de facto* land use practices ongoing across the area of ILUMA WMA, assess how well the WMA was functioning from a wildlife conservation viewpoint, and glean useful insights to inform improved WMA management going forward.

## 2 MATERIALS AND METHODS

### 2.1 Study Area

The study area is located both inside ILUMA Wildlife Management Area (ILUMA WMA), and outside it in the villages immediately surrounding its borders to the west, and in NNP to the east. ILUMA WMA is located in the Kilombero Valley [08°40’S 36°10’E], spanning both Kilombero and Ulanga districts in the Morogoro region of Southern Tanzania (Figure 1). Covering an area of over 7, 000km^2^, the Kilombero Valley is the largest lowland freshwater wetland in East Africa and an established RAMSAR site. ILUMA WMA is nested within this valley, covering an area of 509km^2^ immediately to the west of NNP (United Republic of Tanzania Ministry for Natural Resources and Tourism, 2023). Located in south-eastern Tanzania, NNP is Africa’s largest national park, encompassing 30,893 km^2^ (“Nyerere National Park,” 2023). It was designated national park status in 2019, having previously formed part of the Selous game reserve. ILUMA’s northern boundary is demarcated by the Kilombero river, while areas to the south, west and north contain expanses of agricultural land and villages.

**Figure 1.**
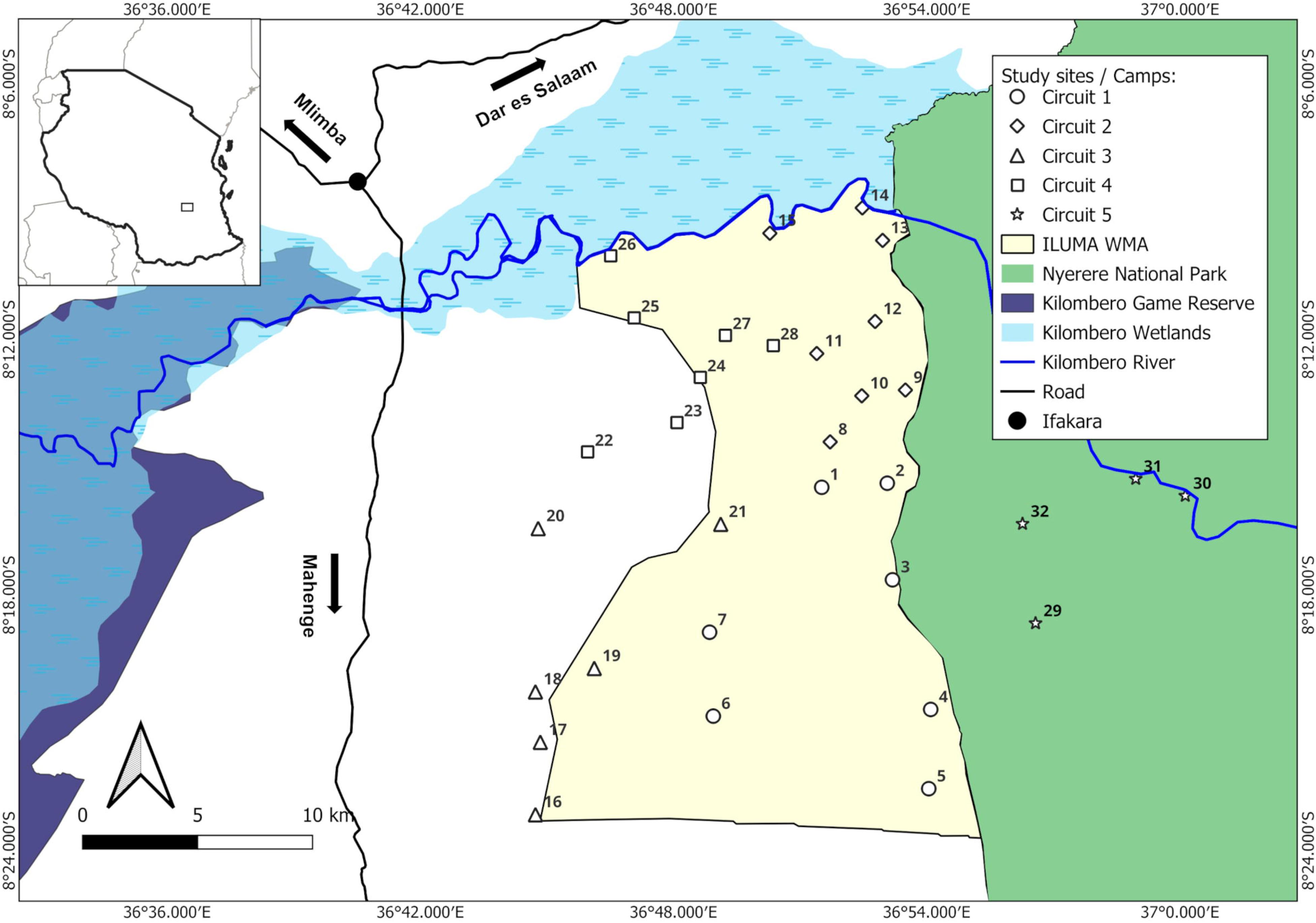
Map displaying the survey area in a national and local context in southern Tanzania. The insert at the top left is a map of Tanzania, while the enlarged area shows the part of the Ifakara-Lupiro-Mangula Wildlife Management Area (ILUMA WMA) to the south of the Kilombero River where most of the surveys were carried out in the context of the nearby Nyerere National Park (NNP) to the east, the extensive wetlands of the Kilombero Valley inland delta upstream to the west and Ifakara town to the north-west.

ILUMA acts as a buffer zone for NNP and includes zones for tourist hunting and local hunting, in addition to a wetland conservation zone. Establishment began in 2011 and user rights were declared in 2015, making ILUMA the 19^th^ WMA in Tanzania (United Republic of Tanzania Ministry of Natural Resources and Tourism, 2015). Land from 14 villages is encompassed by ILUMA (Daconto et al., 2018), and its cover consists of miombo woodlands, dense groundwater forest, riparian forest, open grasslands and wetlands. The climate of the region is typical of moist tropical savannahs with distinct seasons. The main rainy season of reasonably consistent heavy precipitation occurs between March and May each year, with the dry season from June through to October generally seeing little to no rainfall, and then followed by a shorter, less predictable season of generally more sporadic rainfall between November and February. Annual average rainfall in the Kilombero valley typically varies between 1,200 and 1,4000mm (Mombo et al., 2011) and temperatures are highest from December to March.

The survey area consisted of 32 distinct locations, referred to as *camps*, each assigned an identity numbers from 1 to 32 (Figure 2 and supplementary file 1). These camp survey locations were distributed throughout the study area: 22 inside the borders of the ILUMA conservation area, 6 outside it in the domesticated areas close to nearby villages and 4 camps inside adjacent parts of NNP. It should be noted that this study was nested within a larger project related to malaria vector mosquito ecology (Kavishe *et al.,* 2024, Walsh, 2024), the objectives of which were the primary determinant of survey location selection. The logistical requirements of this study therefore had to be adapted to fit within the data collection requirements of that larger multi-disciplinary project. The most important requirement considered when choosing survey sites was year-round availability of surface water, where malaria vector mosquitoes could breed and the survey team could obtain sufficient water for drinking, cooking and bathing (Kavishe *et al*., forthcoming and Walsh *et al.,* forthcoming).

**Figure 2.**
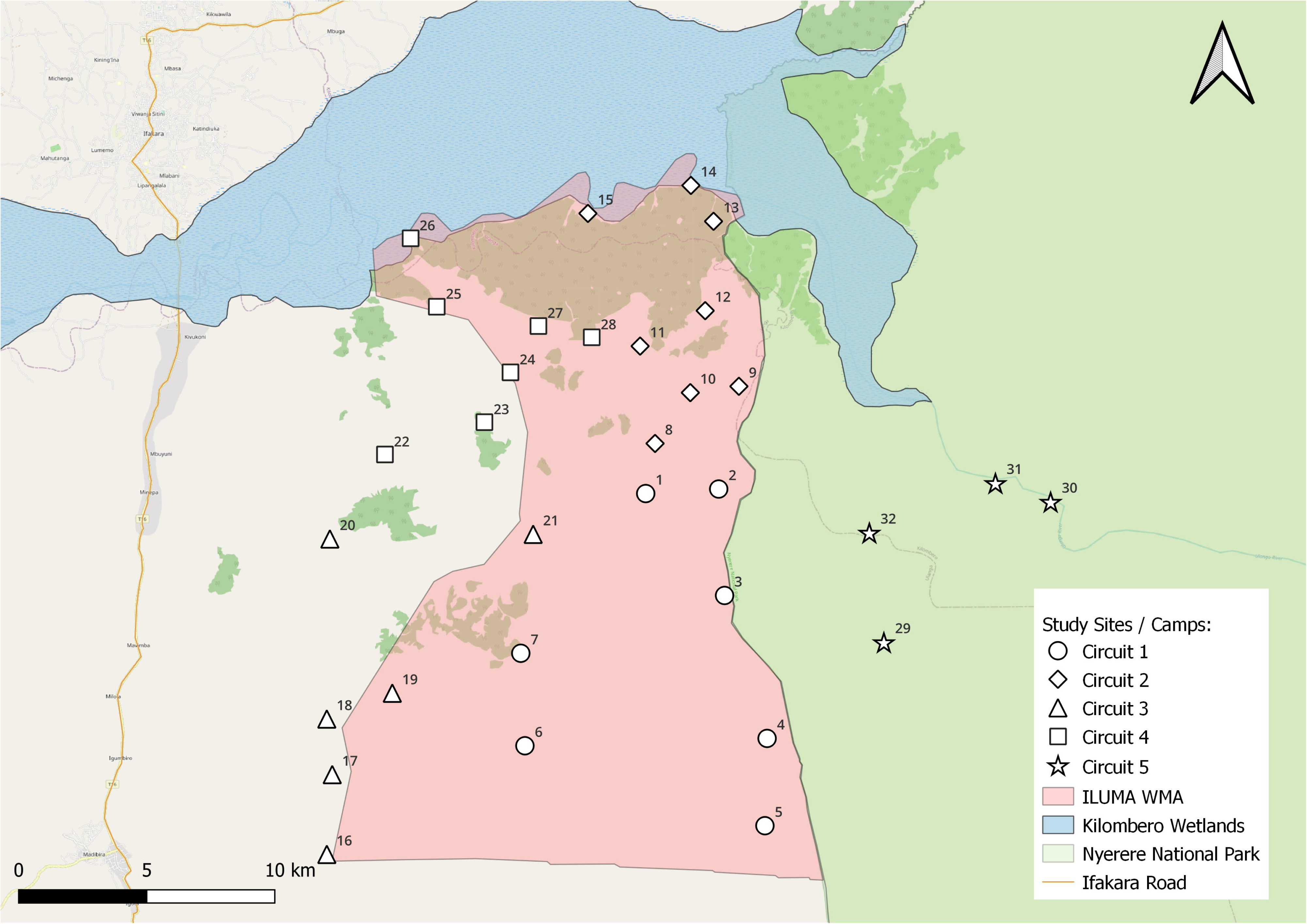
Map displaying the distribution of suitable camping locations used as the sampling frame for all the surveys described in this report. Each of the 32 camps detailed in supplementary file 1 are illustrated in the geographic context of the boundaries of the Ifakara-Lupiro-Mangula Wildlife Management Area (ILUMA WMA) and Nyerere National Park (NNP).

### 2.2 Study design and sampling frame

As explained in detail elsewhere (Duggan, 2024) this study was carried out using a repeated rolling cross-sectional design with three rounds of surveys over the course of one calendar year, specifically 2022. Round 1 ran from 21^st^ January to 16^th^ March, while round 2 ran from 22^nd^ March to 25^th^ May, and round 3 then ran from 25^th^ August to 29^th^ November. Rounds one and two occurred during the wet season and round 3 during the dry season. Each round encompassed most of these 32 camps, but not all of them (Supplementary file 1) for various practical reasons, including physical inaccessibility during the rains or lack of water in the dry season. These camps were assigned to one of five circuits, with each one surveyed over a period of approximately two weeks, staying at each camp for two nights before departing for the next camp the following morning. The radial surveys of various detected human, livestock and wild mammal activities around surface water bodies within a 2km radius of each camp were carried out on the mornings after arriving at each camp (Duggan, 2024, Duggan et al. (2024), forthcoming, Duggan et al., forthcoming).

### 2.3 Formal quantitative radial surveys of human, livestock and wild mammal activities around surface water bodies

As explained in detail elsewhere (Duggan, 2024, Duggan et al. (2024), Duggan et al., forthcoming), radial surveys for various signs of activity by humans, livestock or wild animals (Supplementary file 3) were carried out along the fringes of water bodies within a 2km radius of each camp, These water bodies were first surveyed for mosquito larvae by the accompanying investigators focusing on malaria vector ecology (Walsh, 2024), so the exact direction taken and waterbodies surveyed was primarily determined by the requirements of those entomological surveys. The waterbodies surveyed included seasonal streambeds and the pools of water within them, puddles and pools outside of streambeds, waterholes, pools of water in rice fields and rivers and any other surface water body that might attract mosquitoes, people or animals, even if they were clearly ephemeral in nature. All direct visual observations and indirect signs of activity by humans, livestock or wild mammals (Supplementary file 3) that were encountered along the routes taken between and around the fringes of surveyed water bodies were recorded using the standardized tools provided in supplementary files 4, 5, 6 and 7). The fully explicit data collected through all these formal quantitative observations are presented in supplementary file 8.

### 2.4 Numerical estimation of conventional indices of wild animal community or whole ecosystem integrity based on formal quantitative data from the radial activity surveys

Estimates for each of the following indices of the integrity of wild animal communities or the whole ecosystem at each camp were calculated as follows based on the data obtained through radial activity surveys (Section 2.3). All the following analyses were carried out using the *R*^®^ open-source software package through the *RStudio*^®^ version 2023.03.0+386 environment. While only data relating to activities by wild mammals was used to calculate scores for the first three indices of wild animal community integrity for each camp, the fourth and final index described below, intended as an indicator of whole ecosystem integrity, included all of the variables available in supplementary file 8. Assigned scores for the each of the four following indices at each camp are detailed in supplementary file 2.

*Species Richness Index (SRI)* for each camp was calculated as the absolute number of species in a given area (Whittaker et al., 2001), based on wildlife activity data only, using the *plyr* package and *richness* command. SRI was calculated as: 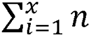, where n = the number of species detected (Moore, 2013), with higher values indicating higher biodiversity.

*Simpson’s Index of Diversity (SID)* were calculated as 1- D, where 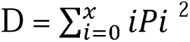, where 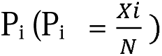 is the proportion of observations accounted for by each species *i*, *x_*i*_* is the total number of each species recorded and N is the total number of individuals recorded.

*Objective Wild Animal Community Integrity Index (OWACII)* was estimated by using the *factoextra* package to carry out a principal component analysis (PCA) of the values for all the various types of detections of all the wildlife species for each camp in the processed datasets provided in supplementary file 8. OWACII represents a synthetic statistical summary parameter, based on this data reduction analysis of all the variables that relate to recorded activities of wild mammals. To make these values easier to interpret, they were normalized by scaling as their Z-score means.

*Objective Natural Ecosystem Integrity Index (ONEII)* was also estimated by PCA using the *factoextra* package (James and McCulloch, 1990; Ortega *et al.,* 2004; Castela *et al.,* 2008; Reza and Abdullah, 2011; Janžekovič and Novak, 2012; Caniani *et al.,* 2016). This synthetic statistical summary parameter represents more comprehensive index of overall ecosystem integrity that OWACII because it is not only based on all the detections of wildlife, livestock and humans but also the recorded summary indicators of land cover at each camp. This index therefore combines all the recorded indicators of wildness with those of encroachment to provide a composite score for comparing the ecological status of different camps. Again, to make these values easier to interpret, they were scaled as their Z-score means.

### 2.5 Cluster analysis to classify survey locations into statistically distinctive categories with common characteristics

Cluster analysis using the k-medoids method was undertaken using the *cluster, dplyr* and *Rtsne* packages, plus *ggplot2* and *scales* for graphing. The silhouette method was used to determine the optimum number of clusters using the *cluster* package. The output suggested two clusters as the optimum, followed by four and then eight (Figure 3). Two clusters were not just the best fit to the data (Figure 3), but also the most practically useful, because the two clusters could be readily and intuitively interpreted as representing degraded versus pristine classes (Figure 4). Plots of Gower dissimilarities were created using *Rtsne* and *ggplot2* packages to reveal clear patterns of clustering.

**Figure 3.**
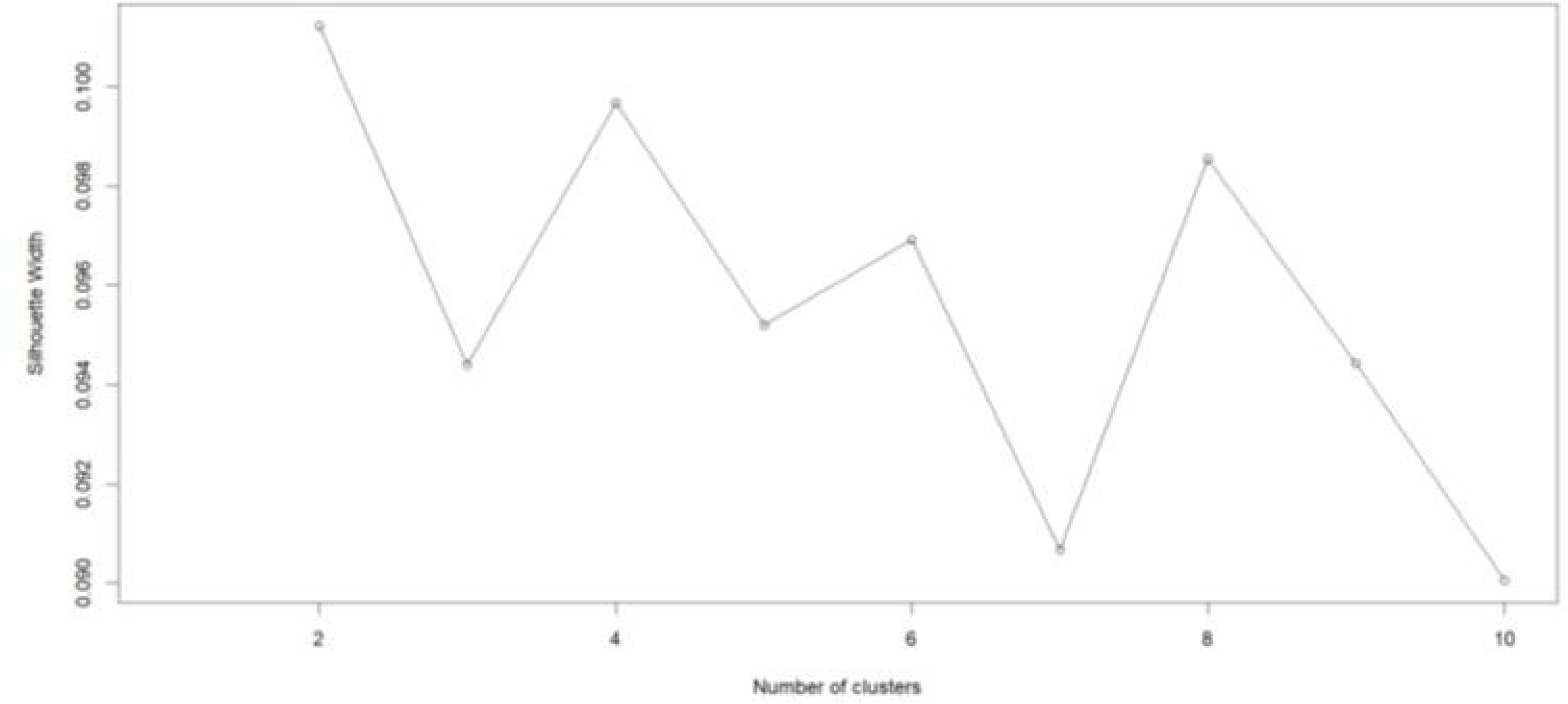
Plot of silhouette widths as output from k-medoids cluster analysis described above. Higher values for width suggest a more accurate number of clusters.

**Figure 4:**
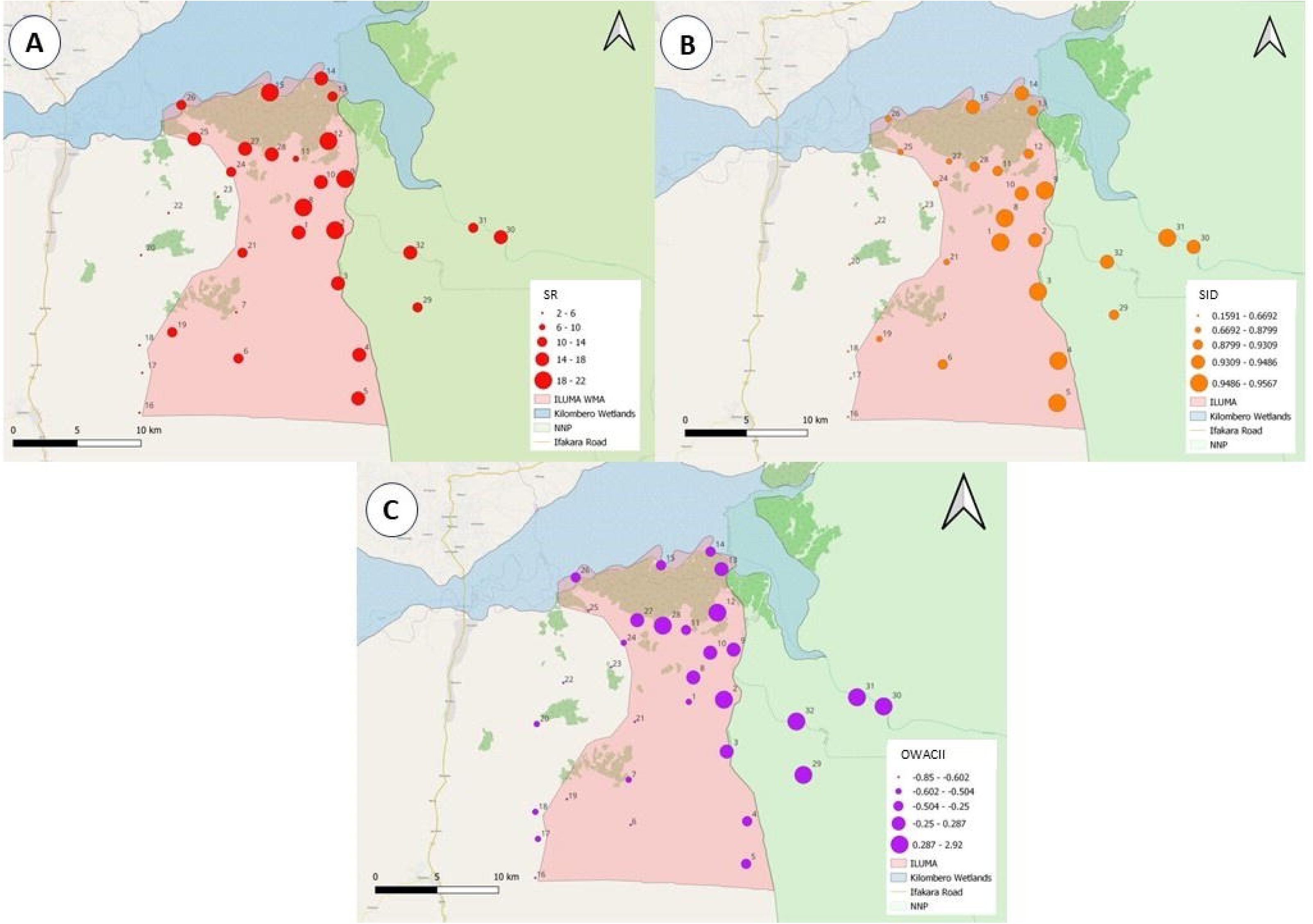
Maps illustrating the distribution of scores for three different indicators of the integrity of the wild animal community across the study area, estimated based on formal quantitative surveys (Sections 2.3 and 2.4) at each of the camps (Section 2.2 and supplementary files 1 and 2): **A**; Species Richness Index (SRI), **B**; Simpson’s Index of Diversity (SID), and **C**; Objective Wild Animal Community Integrity Index (OWACII).

### 2.6 Consensus perception-based estimation of a *Subjective Natural Ecosystem Integrity Index* (SNEII)

Following the three rounds of data collection and prior to any analyses, a *subjective natural ecosystem integrity index* (SNEII) was devised, as an alternative means of assessing the ecological intactness of the area immediately around each camp and the level of degradation, if any, exhibited over the course of the study. This index was scaled as a percentage score, with a score of zero representing a fully domesticated landscape with well-established human settlements and comprehensive land conversion for agricultural purposes, whereas a score of 100% represents a fully pristine, natural ecosystem, with no evidence of human exploitation of any kind.

As described in detail elsewhere (Duggan, 2024, Duggan et al. (2024), Duggan et al., forthcoming) individual scores were carefully assigned based on a consensus reached through discussions among the three investigators on the field team (LD, KW and LT), based on their subjective recollections. The primary and most important criterion for assigning an ecosystem integrity score was the subjective impressions of the intensity of land degradation caused by the conversion of land for agriculture, deforestation for the purposes of charcoal production, timber harvesting or human settlement and for livestock herding. The intensity of degradation of the land surrounding a 2km radius of the camp was determined by estimating the proportion of the natural ecosystem remaining more-or-less intact, based on informal personal observations that were made while present in the camp and during camp radial surveys. This subjectively estimated proportion of intact land was then used as a reference point for initiating the discussions that concluded with the assignment of the score. Camps that evidently belonged on the extreme ends of the scale were assigned first, to provide clear reference points, before those that were less clear and required a more in depth-discussion. The investigators recollections regarding the abundance of wildlife and humans as indicators of ecosystem integrity featured less in conversation when assigning scores but were nevertheless useful as secondary factors to be considered following the discussion of the intensity of land degradation.

These SNEII scores were initially assigned, then discussed in depth, refined and finalized based on consensus (Supplementary file 2) in December 2022. Note that this consensus-based score assignation was completed before the statistical analyses of the numerical summaries of formal, quantitative survey data described above were initiated, to avoid biasing the former towards the latter.

### 2.7 General linear modelling (GLM) analyses

In order to determine which human activities had the most severe impact on natural ecosystems, a series of separate general linear modelling (GLM) analyses were undertaken, in which each of the individual indices described above were treated as an alternative dependent variable, thus acting as one of several potential indicators of natural ecosystem integrity. One exception is that GLM analysis was not carried out on ONEII as variables relating to human activity were used in creating this index, so testing for their association with this derived index would be confounded by circular logic. Each human activity was firstly scaled using Z-score means, using the same calculation method described above. GLM analysis was then carried out in *RStudio*® using the *lme4* package.

A series of univariate GLMs were fitted defining the index as the dependent variable and one given human activity as the sole independent variable. Multiple distributions and link functions were specified until the best fit model with the lowest AIC score was identified. The selected best-fit distribution for SRI was Gaussian family with an inverse link. The selected best-fit distribution for SID was Gamma family with a log link function. The Gaussian family without any link function was chosen as the best fit for both SNEII and OWACII. Multiple univariate GLM analyses were run using each human activity as the independent variable. Then multivariate analyses were then carried out, using a forward stepwise selection procedure to build the most informative and parsimonious explanatory model possible, beginning with the human activity variable with the lowest P value in univariate analysis, which was livestock herding in all cases. Human activities were added and removed from the model based on objective criteria specified *a priori*, until the best fit model based on AIC score was produced: each retained variable was required to at least approach significance in the model it was included in and significantly improve the goodness of fit relative to the otherwise equivalent model from which it was excluded.

## 3 RESULTS

### 3.1 Geographic variations in wildlife activity and ecosystem integrity across the study area

Overall, the study area exhibited a consistent gradient of landscape characteristics, ranging from fully domesticated lands with minimal wild fauna in and around human settlements in the west and south, all the way through to completely intact natural ecosystems with their full complement of wildlife to the north and east (Figure 4 and 5). The overall pattern was readily explained in terms of the geographic distribution of various human activities, even including the establishment of substantial human settlements, which extended deep into the ILUMA conservation area from the villages immediately outside it to the east and south (Supplementary file 9). However, much of the northeast of the WMA was well conserved and even surpassed the rigorously protected survey locations inside NNP to the east in terms of mammalian species diversity. The transition zone between these two extremes inside the WMA exhibited a mosaic of mixed land cover, where wildlife, humans and livestock intermingled.

**Figure 5:**
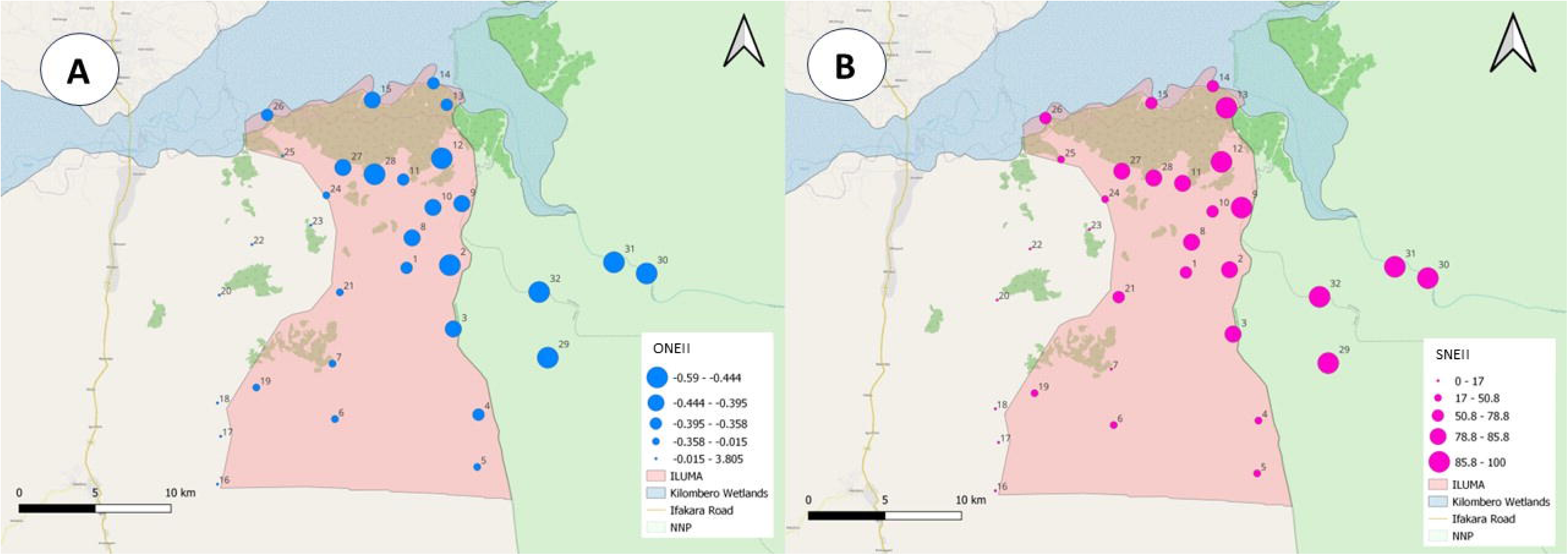
Maps illustrating the distribution of scores for three different indicators of the integrity of the wild animal community across the study area, respectively estimated based on either formal quantitative surveys (Sections 2.3 and 2.4) or subjective appraisal based on consensus investigator perspective (Section 2.5) at each of the camps (Section 2.2 and supplementary files 1 and 2): **A**; Objective Natural Ecosystem Integrity Index (ONEII), **B**; Subjective Natural Ecosystem Integrity Index (SNEII).

The number of wild mammalian species recorded at each camp varied considerably, ranging from 2 (camp number 20 – *Makingi*) to 22 (camp number 12 – *Bwawa la Maya* and camp number 15 – *Mdalangwila*) (Supplementary file 2). Interestingly, all of the five highest ranked camps for SRI occurred in the WMA rather than in the adjacent national park (Figure 4A and Supplementary file 2). This is likely due to the natural cover of the WMA consisting largely of miombo woodland and groundwater forest habitats which appears to host to a larger diversity of species than the acacia savanna habitat that dominates most of the camps sampled in NNP.

The range of SID values calculated for each of these camps for this index varied widely (Figure 4B ad supplementary file 2). The most diverse camp *Bwawa la Nyati* had a score of 0.96, while the least diverse camp, *Mavimba Porini* scored 0.16. Exactly half of the camps have a SID score >0.90, which confirms that the better conserved areas of the WMA are highly species diverse. Of the top five camps all are located inside ILUMA in locations with Miombo woodland land cover. While the camps located inside NNP did yield high SID scores (all > 0.90), none of them were ranked in the top 5. This observation illustrates how even a degraded miombo woodland ecosystem can support a greater diversity of species than a well-protected, non-encroached acacia savanna.

However, the top ranked camps in terms of OWACII were the four NNP camps (Figure 4C and supplementary file 2). Meat poaching was only ever recorded at one of these locations inside NNP and neither cattle herding nor any of the other observed human activities were recorded at any of them (Supplementary file 9). The five worst-ranked camps in terms of OWACII all contained large areas of human settlement where extensive human activities were ongoing. Similarly, the top ten scoring camps in terms of ONEII were all located either inside NNP (all ranked in the top 4) or in the northeastern section of ILUMA, very close to the border of the national park, while those with the lowest scores were all either in the neighbouring villages to the west or around heavily settled areas inside the eastern and southern boundary of the WMA. Overall, the SNEII estimates were closely correlated with both of these objective alternatives, ONEII in particular (Duggan, 2024, Duggan et al. (A), forthcoming, Duggan et al. (B), forthcoming), so the cartographic picture obtained with this subjective approach was very similar (Figure 5), even though it was based on consensus investigator perceptions rather than statistical synthesis of formal quantitative activity surveys.

### 3.2 Influences of specific human activities on wildlife and ecosystem integrity

For all three of the objective indices of wild animal community diversity (SRI, SID) and integrity (OWACII), single independent variable GLM analyses indicated that a number of human activities were associated with variation of the index in question, but only a small subset proved significant and were retained in the subsequent final multiple independent explanatory variable analysis (Supplementary file 10). Livestock herding was the only human activity to significantly impact species richness in multivariate analyses of SR. In multivariate analysis of SID, however, livestock herding and unauthorized fishing were the only two significant human activities. Both of these variables were associated with lower SID scores, but the effect size for the former was more than twice the latter. Livestock herding and meat poaching both appeared associated with lower ONACII scores in multivariate analysis, each having an apparently a similar magnitude of impact.

While similar GLM analyses could not be applied to assess the influence of these activities upon ONEII, because this dependent variable was estimated using these same activities that would also be treated as the independent variables (Section 2.6), multiple independent explanatory variables analysis revealed that six human activities were predictive of the SNEII (Table 1). It therefore appears that the perception based SNEII represents a far more sensitive, statistically powerful dependent variable with which to elucidate the associations of ecosystem conservation successes and failures with various human activities than any of the more objective alternatives based on formal quantitative surveys of wild animal activity (Duggan, 2024, Duggan et al. (B), forthcoming). Livestock herding, charcoal burning, timber harvesting, human settlement and both rice and other tillage agriculture all proved significant predictors of SNEII in the multivariate analysis presented in table 1, so these relationships are represented cartographically in supplementary file 11.

**Table 1.** Results of single and multiple explanatory variables generalized linear modelling analyses of scaled human activity indicators as determinants of the estimated Subjective Natural Ecosystem Integrity Index (SNEII) for each camp surveyed, with statistically significant associations highlighted in bold. The best fit models identified for this SNEII outcome all assumed a Gaussian distribution and identity link function for the dependent variable.

| Activity | <u>Single Independent Variable</u> | <u>Multiple Independent Variables</u> |
| --- | --- | --- |

| | $\beta \pm \text{SEM}$ | T | P | $\beta \pm \text{SEM}$ | t | P |
| --- | --- | --- | --- | --- | --- | --- |
| Livestock Herding | **-29.3 ± 3.1** | **-9.51** | **<<0.0001** | **-17.0 ± 2.9** | **-5.85** | **<<0.0001<sup>a</sup>** |
| Charcoal Burning | **-14.5 ± 5.6** | **-2.59** | **0.0146** | **6.8 ± 2.7** | **3.01** | **0.0058<sup>a</sup>** |
| Timber Harvesting | -7.7 ± 6.0 | -1.27 | 0.213 | **4.4 ± 2.0** | **2.25** | **0.0333<sup>a</sup>** |
| Fishing | 1.1 ± 6.2 | 0.18 | 0.862 | 3.6 ± 2.4 | 1.50 | 0.1454<sup>b</sup> |
| Hunting | -4.9 ± 6.1 | -0.80 | 0.429 | -0.88 ± 2.2 | -0.40 | 0.6917<sup>b</sup> |
| Human Settlement | **-23.3 ± 4.5** | **-5.19** | **<<0.0001** | **-6.5 ± 2.7** | **-2.38** | **0.0256<sup>a</sup>** |
| Well | **-21.4 ± 4.8** | **-4.49** | **0.0001** | 3.3 ± 3.0 | 1.10 | 0.2814<sup>b</sup> |
| Meat Poaching | -8.2 ± 6.0 | -1.37 | 0.180 | -1.2 ± 2.0 | -0.62 | 0.5388<sup>b</sup> |
| Human Presence | **-21.6 ± 4.7** | **-4.6** | **0.0001** | 5.4 ± 3.2 | 1.70 | 0.1011<sup>b</sup> |
| Rice Farming | **-27.9 ± 3.5** | **-8.00** | **<<0.0001** | **-9.4 ± 2.9** | **-3.25** | **0.0033<sup>a</sup>** |
| Other Tillage Farming | **-29.4 ± 3.0** | **-9.69** | **<<0.0001** | **-9.7 ± 3.7** | **-2.59** | **0.0157<sup>a</sup>** |

SEM: Standard error of the mean for the $\beta$ coefficient estimate.
<sup>a</sup> As estimated from the final, most parsimonious best-fit GLM, in which this was included on the basis of either being a significant ( $P \leq 0.05$ ) determinant of SNEII or approached significance ( $P \leq 0.10$ ) as a determinant of SNEII and significantly ( $P \leq 0.05$ ) improved the goodness of fit based on the estimated Akaike Information Criterion (AIC).
<sup>b</sup> As estimated from the last GLM in which this variable was still included before being removed on the basis that it was not a significant determinant of the SNEII ( $P > 0.05$ ) or
approached significance ( $P \leq 0.10$ ) but did not significantly improve the goodness of fit based on the estimated AIC ( $P > 0.05$ ).

While charcoal burning and timber harvesting led to an increased SNEII score in multiple independent variables, this is most likely due to reverse causality: These activities obviously do not improve ecosystem integrity but rather tended to take place in areas inside the WMA with moderate to high SNEII scores, where the prerequisite natural resources remained available (Supplementary file 11, figures S11.2 and S11.3).

Interestingly, cattle grazing was recorded just as frequently at the degraded camps inside ILUMA as at those in fully domesticated areas outside the WMA (Supplementary file 11, figure S11.1), where this activity is not restricted but must be traded off against agricultural land use options during the growing season for obvious reasons. It is also notable that cattle herding was encountered remarkably frequently at some relatively pristine locations with moderate to high SNEII scores inside the WMA (Supplementary file 11, figures S11.1), suggesting that grazing *per se* may not, in itself, have as great an impact as that suggested by table 1. Having said that, one evident trend was that livestock herding had a consistently significant negative impact on all the estimated indices of wild animal community (Supplementary files 10 and 11) and overall ecosystem integrity (Table 1) and was the single most quantitatively important variable in all four multivariate models presented in table 1. Furthermore, no other human activity was consistently associated with all of these indices in multivariate analyses (Table 1, supplementary files 10 and 11).

In contrast, human settlement appeared the least influential of the four activities found to be associated with lower SNEII scores in the multivariate analysis detailed in table 1: Once the effects of the following other activities was accounted for, the effect size for human settlement appeared similar to or less than that for farming of rice or other tillage crops and far less than that for cattle herding (Table 1). Furthermore, it was not significantly associated with reductions of any of the indicators of a healthy wild animal community detailed in supplementary file 11. One obvious reason for this surprisingly modest and inconsistent relationship between ecosystem integrity and human settlements was that the 3 northernmost camps inside the WMA, namely camps 14, 25 and 26 along the banks of the Kilombero River counterintuitively had both substantial human settlements and remarkably intact natural land cover (Figure 6).

**Figure 6:**
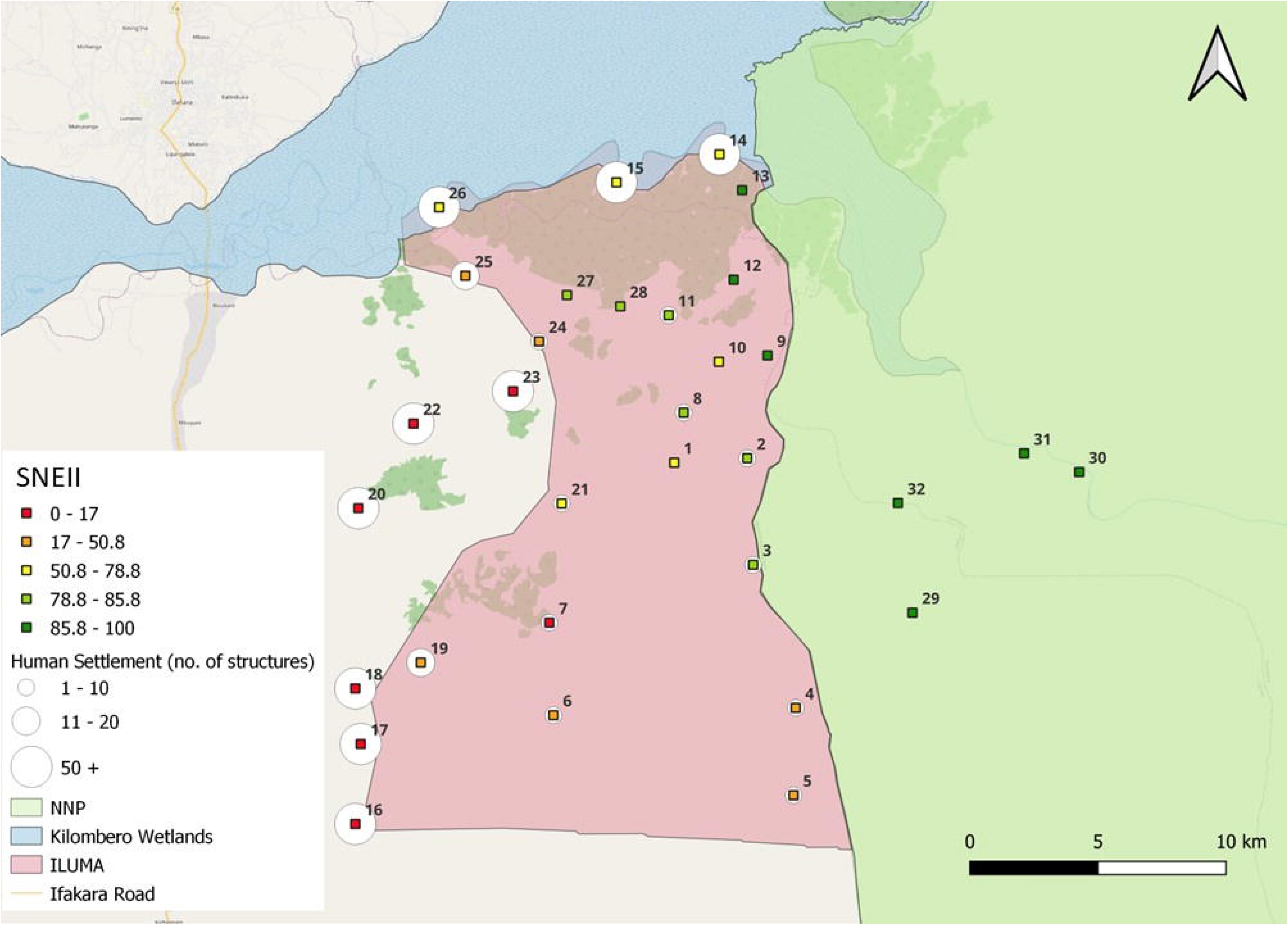
Map of ILUMA WMA showing each camp location’s Subjective Natural Ecosystem Integrity Index (SNEII) score, ranging from the lowest (red) to the highest (green) scores, and layered over the extent of human settlement estimated for each camp, because the latter was identified as a significant predictor of the former in table 1.

### 3.3 Cluster analysis: General rules and informative exceptions

Experience-informed interpretation of a cluster analysis of survey location characteristics revealed that these three planned, carefully managed human settlements within the WMA actually appeared beneficial in conservation terms. Cluster analysis split the 32 camp locations into two equally sized, clearly distinguished groups composed of 16 camps in each (Supplementary file 2), which were clearly and intuitively related to their geographic distribution (Figure 7) and level of environmental conservation or degradation (Figures 4, 5, 6, and 8 plus supplementary files 2, 9, 10, 11 and 12).

**Figure 7.**
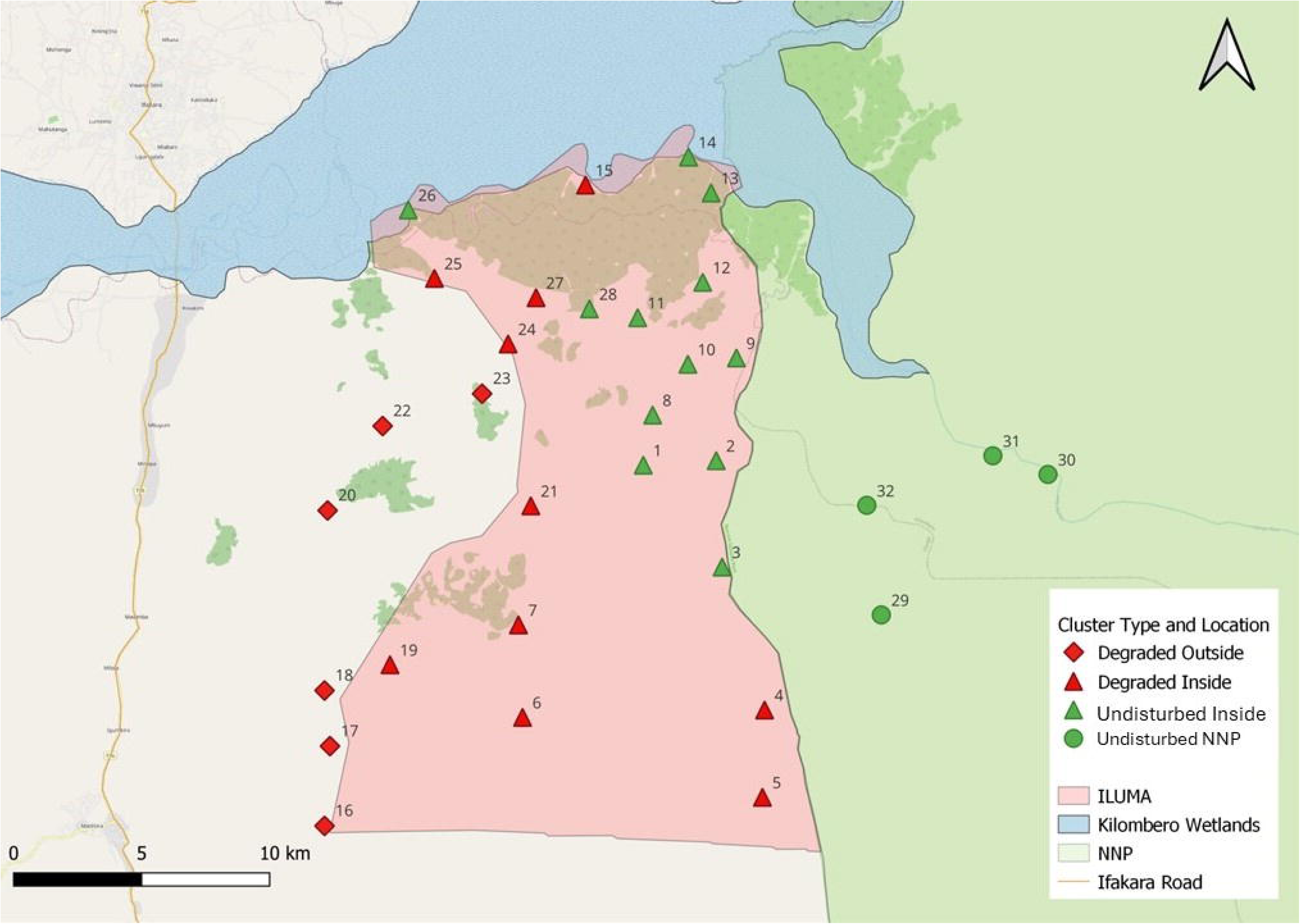
Map of ILUMA WMA displaying the 32 camp locations sampled and the cluster they were assigned to by k-medoids analysis as described in Section 2.5. Shape represents location, either inside ILUMA WMA, outside in the villages along its western boundary or inside NNP to the east, while colour represents assignment to either the *pristine* (green) or *degraded* (red) cluster, the interpretations of which based on figures 4, 5, 6 and 7, as well as supplementary files 2, 9, 10, 11 and 12).

The first cluster contained all of the village-based camps outside the western border of ILUMA that were fully domesticated, as well as highly encroached camps inside ILUMA to the west and to the south with similarly *degraded* land cover. In stark contrast, all the camps statistically attributed to the second cluster were better conserved, so this cluster was interpreted as reflecting a relatively intact natural landscape with natural vegetation cover and wild fauna. These *pristine* camps occur where ILUMA borders NNP on the eastern side of ILUMA. where there is less encroachment by humans and livestock, and to the north where the Kilombero river acts as a physical boundary against encroachment. This second cluster also included the NNP camps to the east.

Generally speaking, the geographic patterns identified by cluster analysis were clear and consistent with those apparent from maps of the various estimated indices of the integrity of the wild animal community or the overall ecosystem (Figures 4, 5, 6 and 7, as well as supplementary files 2, 9, 10, 11 and 12). There were no pristine camps outside the western boundary of ILUMA and there are no degraded camps inside NNP. ILUMA contains a mix of both degraded and pristine, but with more degraded camps located to the west and south of the area and more pristine camps located to the east and north. Broadly speaking, this overall pattern mirrored the geographic distribution of all the various human activities observed, notably the establishment of settlements that were typically surrounded by a range of other unauthorized activities (Figure 9A).

**Figure 8.**
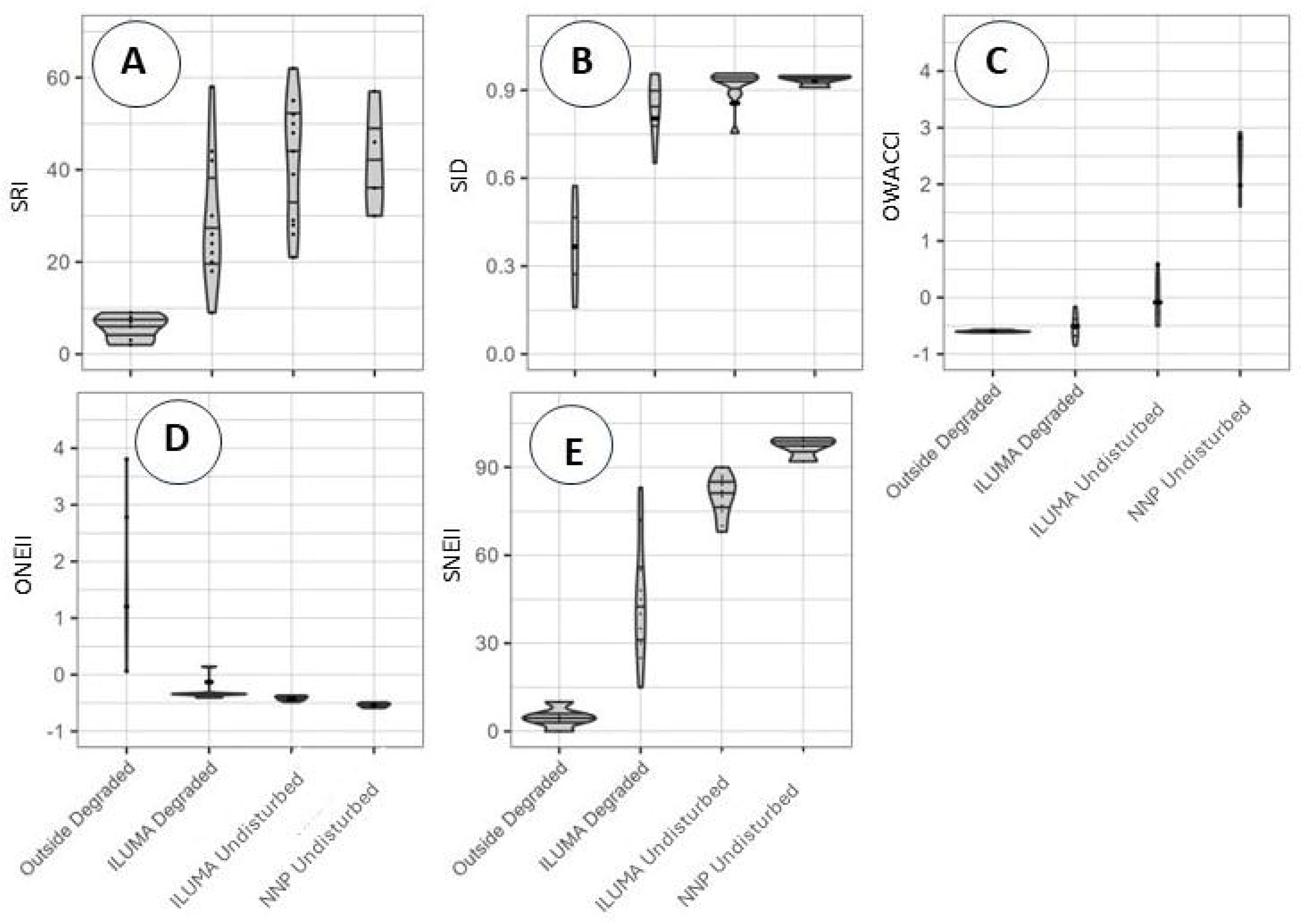
Dot and violin plots of the estimated values for all of the various indices of wild animal community and overall ecosystem integrity (Supplementary file 2) for each camp classified by location and cluster assignment as illustrated in figure 4: **A**; Species richness index (SRI), **B**; Simpson’s index of biodiversity (SID), **C**; Objective wild animal community integrity index (OWACII), **D**; Objective natural ecosystem integrity index (ONEII) and **E**; Subjective natural ecosystem integrity index (SNEII).

**Figure 9:**
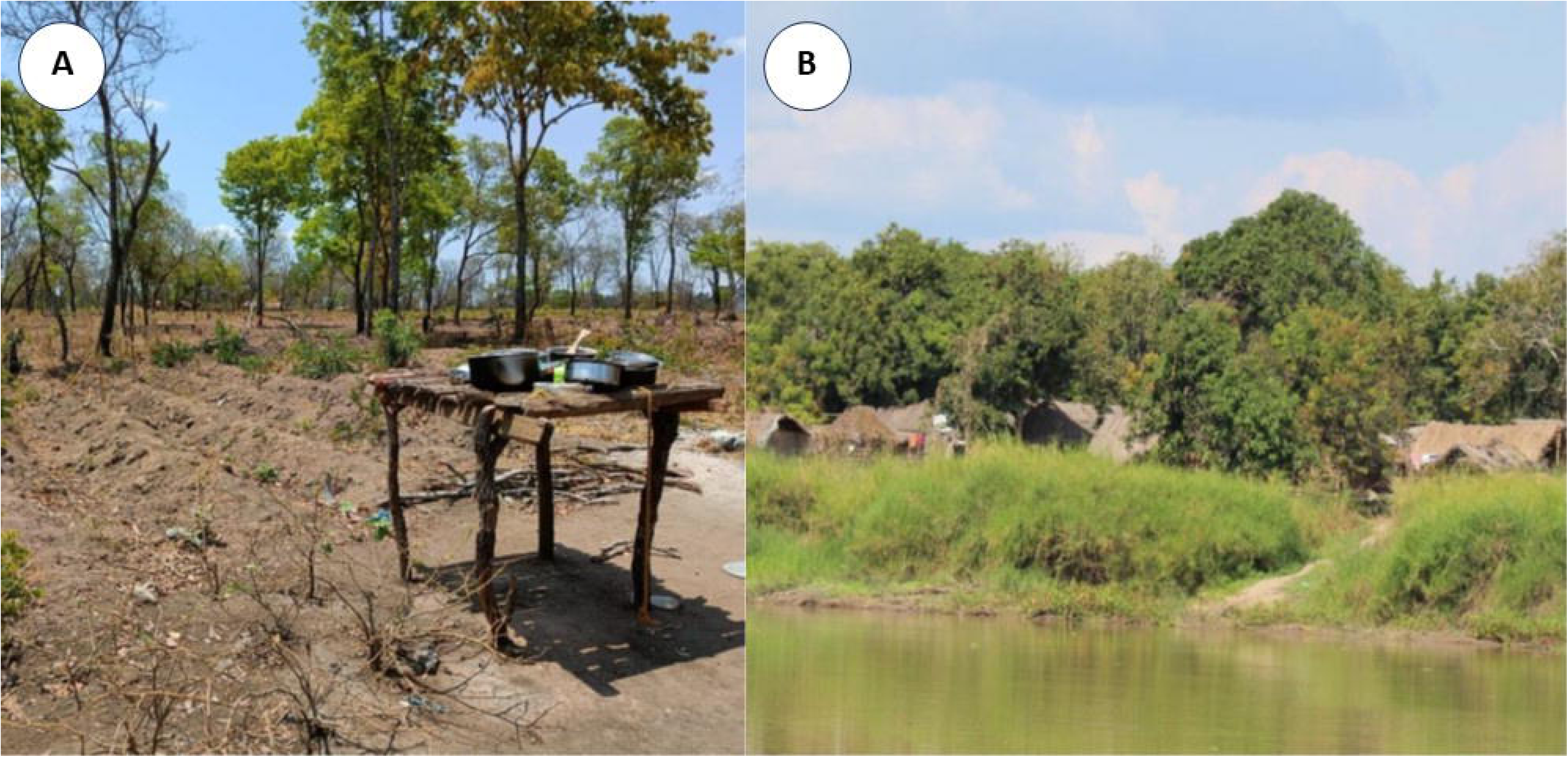
Photographic illustrations of how human settlements inside the ILUMA WMA were usually observed to be surrounded by comprehensive degradation of the natural ecosystem (Panel **A**) but were also noted to have negligible impact upon the surrounding natural terrestrial ecosystem in the case of some authorized fishing camps established along the Kilombero River (Panel B). Panel **A** depicts a highly encroached area visited during a radial survey around camp 4, with obvious evidence of land conversion from forest to tillage agriculture and human settlement. The image in panel **B** was taken at camp 14, depicting how this authorised fishing settlement on the banks of the Kilombero river had negligible impact upon the groundwater forest immediately behind it. Note the contrast in level of land clearance between these two settlements.

However, consistent with figure 6 some surveyed locations within the ILUMA conservation area that had authorised fishing camps (camps 14. 15 and 26) as part of the agreed management plan for the WMA were assigned to the pristine cluster by the objective statistical cluster analysis (Figure 7). These three authorized fishing villages established within the WMA conservation area therefore represented informative exceptions to the general rule that human settlements, and all the various human activities that are otherwise closely associated with them, appear inevitably associated with habitat degradation (Figures 6 and 9A). These fishing camps were well managed settlements that were confined to a very small geographical area, with an authorised and carefully regulated sustainable livelihood (fishing) that has negligible impact on the adjacent terrestrial habitat (Figures 6 and 9B). Consequently, the surrounding forest and floodplain were observed to be essentially intact with no sprawl of dwellings beyond the designated settlement limits, and experienced minimal encroachment due to being monitored and patrolled frequently (Figures 6 and 9B).

## 4 DISCUSSION

Data detected, gathered, and analysed throughout the course of this study indicate that there was both adherence to, and at times strong deviation from, the land use plans that had been negotiated and agreed with the stakeholder communities for ILUMA WMA. Unauthorised human activities were recorded most frequently close to the western and southern boundaries of ILUMA, where the WMA conservation area is bordered by formally established villages. As a general trend, the southwest experienced the highest levels of unauthorised human activity, which lessened in frequency approaching the northeast of the WMA and the eastern boundary with NNP.

Unauthorised human settlement marks a clear breech of the agreed land use plans, and the extractive and ancillary activities that are associated with human settlement such as land clearing or deforestation for building materials and agriculture, all alter the landscape and ecosystem functioning. Most unauthorised human settlements within ILUMA’s boundaries were clearly associated with high levels of land clearing and extractive activities. On the African continent, the area south of the equator accounts for 13% of the land globally that is facing degradation due to conversion of natural lands for agriculture or encroachment into protected areas, a phenomenon which is increasing in both extent and severity across the earth (Bai et al., 2008). This degradation of natural ecosystems on a global scale is a crucial factor in why it is imperative for small-scale community-based conservation programmes, such as the one in place here, to be well maintained and in doing so could prove vital in preventing land degradation.

Rice farming took place on a much large scale throughout ILUMA. Rice agriculture is primarily a cash crop in the Kilombero valley, grown largely for sale rather than subsistence (Nkuba et al., 2016), so it is correspondingly grown wherever suitable soil allows. The practice of rice farming in the Kilombero has long been established due to the largely flat topography of this low-lying valley that seasonally floods and fertile *fluvisol* soil type that can retain water for long periods. Due to the nature and practice of rice paddy growth, and the large amount of water it requires, in many areas where rice farming was recorded in ILUMA degradation of a waterbody nearby was evident. The majority of rice cultivation in ILUMA involved some level of water course diversion for irrigation, whether from ephemeral waterholes in the dry season or streams. The diversion of water from its original path caused waterholes to dry out quicker and provided less resources for the wildlife that depend on them. A study on the Kilombero wetlands, part of which was included in this study area, found that the flow of many rivers has been reduced and that a number of swamp areas have been reduced in size or dried up completely as a result of intensive agriculture and deforestation in the Kilombero wetlands (Mombo et al., 2011).

Overall, while livestock herding has appeared to be the most statistically obvious factor directly associated with to species richness, diversity, and ecological integrity in ILUMA WMA over the course of this study, it was also the most frequently detected unauthorised human activity. Indeed, the most frequently detected unauthorised human activity in ILUMA was undoubtedly livestock grazing, with over 1400 instances cattle herding recorded. It should therefore be cautioned that other unauthorised human activities, like cultivation of rice and other tillage crops, may have at least comparable impacts that might not necessarily be as readily attributable to them in statistical terms.

Cattle grazing was encountered just as often around the degraded camps inside ILUMA as around those outside it, where herding is not restricted but has to compete with agriculture as a land use option. Although it is clear in the land use agreement (District Authorities, 2016) that cattle grazing is not a permitted activity inside of ILUMA’s boundaries, the evidence reported herein illustrates just how extensively this aspect of the WMA plan has been breached. Indeed, the informal observations and anecdotal evidence accumulated by the investigators indicate that grazing pressure on lands around the ILUMA WMA was being alleviated by unauthorised cattle herding within the supposedly protected area, especially close to its the western and southern boundaries.

ILUMA’s habitat type is predominantly miombo woodland of different densities, and cattle herding in Tanzanian miombo woodland, in an area with similar abiotic and biotic characteristics to that assessed in this study, revealed mixed ecosystem responses to cattle grazing (Mtimbanjayo and Sangeda, 2018). High intensity cattle grazing caused a significant negative effect on both the density and diversity of miombo tree species, while less intense herding, especially during the dry season, was shown to have no adverse effect on the ecosystem (Mtimbanjayo and Sangeda, 2018). Given the somewhat flexible nature of land use agreements for WMAs in Tanzania, regulated seasonal grazing is permitted in some WMAs, such as the Enduimet WMA in the north of the country (University of Copenhagen Department of Food and Resource Economics, 2015). It may therefore be useful to consider whether rigorously regulated seasonal grazing in ILUMA might help to control the level of unregulated herding, by permitting it at certain times and in certain areas only, and in doing so protecting the areas that are highly valuable for wildlife. The village lands outside ILUMA are largely used for the production of agricultural crops during the wet season, thus displacing cattle into the nearby WMA. Seasonally allowing limited cattle grazing inside the WMA during the rains might not only allow more controlled regulation of this important local livelihood, it might also enhance productivity of this protected area by preventing the overgrowth of perennial grasses and encouraging shorter more-nutritious grasses to grow, thus benefiting both livestock and wildlife (Fynn et al., 2016). Indeed, it is notable that cattle grazing was common at some locations at the interface between relatively intact and heavily degraded lands with intermediate to high scores for SRI, SID, OWACII (Supplementary file 10), ONEII and SNEII (Supplementary file 11, figures S11.1), suggesting that this very common land use activity may not necessarily have as large a negative impact upon wildlife populations and ecosystem integrity as face value interpretation of table 1 might otherwise indicate.

Limiting regulated grazing within the ILUMA WMA to the wet season may also be remarkably compatible with wildlife migration patterns and complementary non-extractive income-generating activities. Several important wild mammalian species, notably zebra, sable and hartebeest are known by the VGS and park rangers to migrate westwards into the open acacia savannah areas of NNP during the rains to access short green grass with high protein content (Estes, 2012) so the grazing of cattle within the WMA during this season, when taller grasses suitable for bulk grazing is most abundant, should minimize competition with wildlife over resources. Conversely, these same wildlife species tend to migrate back into the ILUMA WMA during the dry season, to exploit its extensive miombo woodland, floodplain grasslands and groundwater forest habitats. The dry season, when livestock may be returned to fallow agricultural land, typically runs from June through October, thus overlapping with the summer holiday season of tourists from Europe and North America. The dry season is therefore an ideal time for hosting tourists, at a time when it is most convenient to exclude cattle grazing. However, based on the results presented herein, it is evident that approximately two thirds of the WMA is already too degraded for tourism and that current rates of livestock encroachment into the WMA are unsustainable. Without any interventions like a rigorously regulated wet season grazing plan, to both regulate this important local livelihood and disincentivise potentially more destructive ones like land conversion for agriculture, continued encroachment may well result in this WMA becoming non-viable as a conservation area.

Fortunately, evidence of success with respect to the established practice of another natural resource utilization activity that conducted in an authorized, carefully regulated manner within the WMA conservation area is encouraging, suggesting that there may be scope for participatory development of such compromise solutions. In stark contrast to all the aforementioned breaches of the ILUMA land use plan across the southern and western parts of the supposedly protected area of the WMA, one section of the management plan that was effectively adhered to over the course of this study was the authorised fishing camps in the north of ILUMA, along the banks of the Kilombero river. Very few unauthorised activities were detected at these locations despite the presence of substantive long-established fishing villages, and the areas immediately outside of these settlements appeared ecologically intact with very little evidence of degradation (Figures 6, 7 and 9B). Crucially, not only do the residents of these locations have permission to extract fish from the Kilombero river and sell it as a means of making a living, they also can and do call upon the WMA management and VGS to enforce fishing regulations and apprehend unlicensed competitors from outside the area.

This vital collaborative relationship between the WMA authority and one of the stakeholder communities it serves was reflected in many obvious ways during surveys conducted in and around these villages. Fishing is carried out with nets or by hook and line, from dugout canoes hollowed from tree trunks and is traditional in its forms of practice. Based on both formal data collected and informal observations made while conducting the field work, notably conversations with the VGS about their interactions with these villages, the residents of these fishing settlements, generally speaking, agreed with and adhered to the WMA the land use plan. All three villages, plus two others along the same stretch of river that were not included in the sampling frame, all maintained human settlement patterns that were confined within a small area. These settlements are also much smaller in size than the formally established villages outside of ILUMA, and no large-scale agriculture was recorded in the vicinity of the three relevant camps. All of these fishing villages were surrounded by fully intact, dense groundwater forest within meters from the centres of these settlements.

The presence of authorised residents may actually be helpful in deterring unlawful activity in the areas surrounding these settlements. Due to the mutually beneficial nature of the fishing camps for both the residents, who practice sustainable livelihoods provided by the intact natural ecosystem that surrounds them, residents are quick to report suspicions of illegal activities within the WMA to the VGS. This evidence clearly demonstrates that well-managed human settlements within a protected area can, not only have minimal direct impact upon on the ecological intactness of an ecosystem, they can actually bolster its conservation through resident community vigilance and reporting on a year-round basis. Such passive but continuous surveillance by resident communities living within a protected area contrasts with, but also complements, more active but intermittent forms of surveillance by community-based professionals like VGS conducting patrols.

Overall, this success story illustrates how sustainable livelihoods for local people and effective ecological conservation may be achieved concurrently if adherence to formally agreed land use plans can be adhered to through compromise and collaboration between participatory conservation authorities and the resident stakeholder communities that live in and around such such protected areas. Learning from this example, an analogous approach to negotiating seasonal grazing agreements with villages adjacent to the WMA, or even those already established within it, might involve conditionally allowing limited use of protected rangelands within the WMA, in exchange for de facto participation of beneficiary communities in collaborative regulation of not only grazing itself, but also potentially more destructive activities like agriculture and extractive deforestation.

Such compromise solutions may prove crucially important to delivering on the roles that WMAs, and other sundry institutional models for community-based management of fringe conservation areas, are meant to play as buffer zones around more vertically managed, rigorously protected parks and reserves. Despite the extensive encroachment seen in the west and south of the ILUMA WMA, the gradient of wildlife activity across the WMA clearly increases moving eastward and northward as the conservation status of the land improves and evidence of illegal human encroachment dissipates, with some parts of the WMA having similar ecosystem integrity to the adjoining national park, not to mention higher mammalian biodiversity. The ILUMA WMA clearly does indeed act as an important buffer zone for NNP by providing a protected area with extensive woodland, forest and floodplain habitats that are useful for many species, including many from inside the national park that use it on a seasonal or opportunistic basis.

The remaining pristine areas of ILUMA offers invaluable habitat supporting a wide diversity of wild mammals, further emphasising the need for creative solutions to achieve collaborative, community-based regulation of human encroachment into the WMA. Beyond merely halting ongoing ecosystem degradation in such compromised fringe conservation area across the tropics, it has never been more important to restore them to their intact natural state (Strassburg, 2020, Wilkinson, 2019) so the ongoing evolution of effective institutional models for their devolved governance and management is of paramount importance. Indeed, more effective schemes for managing vulnerable forested areas in the face of growing human population pressure have a particularly important role to play in simultaneously combating climate change (Strassburg, 2020) and the emergence of novel pathogens capable of causing pandemics (Wilkinson, 2019). The natural land cover of ILUMA is predominantly miombo woodland - an ecosystem type with deep soils, abundant hardwood trees and very high carbon sequestration potential when it recovers: The wood of regrowing miombo trees sequesters 0.5 to 0.9 tons of carbon ha^-1^per year, while soils in a regrowing miombo woodland sequester >100 tons of carbon ha^-1^per year (Williams et al., 2008). Furthermore, the biodiversity of mammals (and presumably the pathogens they carry) was highest in parts of the study area with miombo woodland vegetation cover, notably in where it overlapped with various human activities locations at the scrubby interface between degraded and pristine land (Supplementary files 1, 2, 10).

## 5 CONCLUSION

Across much of ILUMA WMA, in areas where agreed land use plans were not adhered to, rampant cattle herding and land clearing for agriculture were associated with reductions in wildlife richness and biodiversity, as well as overall ecosystem integrity. Although human settlement was also generally associated with reduced natural ecosystem integrity, some notable exceptions illustrate how sustainable livelihoods for local people that are based on well-managed natural resource harvesting practices may actually enhance conservation effectiveness: Several authorised fishing villages within the WMA, where fishing was permitted and local communities collaborated with the WMA management to enforce conservation regulations, were surrounded by pristine land cover with thriving terrestrial wildlife populations. Correspondingly, the best conserved parts of the WMA not only included those closest to the boundary with the national park to the east, but also these fishing villages along the riverbank to the north, where compliance with agreed land use plans was most rigorous. Applying lessons learned from this success story more broadly across the WMA, the offer to negotiate conditional grazing rights within the ILUMA during the rainy season, when cattle need to be moved off neighbouring agricultural land, might motivate beneficiary communities to participate in more effective collaborative interventions against unsustainable levels of ongoing encroachment. Overall, this study illustrates how well-managed WMAs may host resident local communities undertaking selective natural resource extraction activities and collaborate with them to achieve effective *de facto* conservation practices, thus maintaining robust wildlife populations and acting as effective buffer zones along the fringes of larger core conservation areas like national parks and game reserves.

## DATA AVAILABILITY STATEMENT

**Supplementary file 1;** Table S1: The number, name, location, coordinates, and ecological characteristics of each camp location, together with the quadrant circuit to which it was assigned and the number of times it was surveyed https://doi.org/10.5281/zenodo.11099575

**Supplementary file 2;** Table S2. Scores for various indices of the integrity of the wild animal community or ecosystem as a whole, estimated for each surveyed camp location (Section 2.2) based on either a consensus approach to synthesizing investigator-perception (SNEII, Section 2.6) or numerical syntheses (SRI, SID, OWACII and ONEII, Section 2.4) or cluster analysis (Section 2.5) of extensive data from formal, quantitative surveys of observable human, livestock and wildlife activities, as well as land cover characteristics (Section 2.3 and Supplementary file 8) https://doi.org/10.5281/zenodo.11099588

**Supplementary File 3;** Figures S3.1 to S3.4: Photographs illustrating different ways in which signs of livestock, humans or wild mammal activity, as well as land cover attributes, were observed and classified https://doi.org/10.5281/zenodo.11099593

**Supplementary File 4;** Complete list of all wild herbivores, wild carnivores, wild primates and prosimians and wild rodents sampled in this study https://doi.org/10.5281/zenodo.11099600

**Supplementary File 5;** Data collection key devised and refined during a preliminary pilot study between October and November 2021 and used throughout the course of the field study to inform accurate and consistent recording of data https://doi.org/10.5281/zenodo.11099602

**Supplementary File 6;** Criteria defining terms used in data dictionary key https://doi.org/10.5281/zenodo.11099616

**Supplementary File 7;** Data collection sheet developed during preliminary investigation in October and November 2021 and used for all data collection in both transect segment surveys and camp radial surveys throughout the course of the study https://doi.org/10.5281/zenodo.11099618

**Supplementary File 8**; The full data set derived from formal quantitative surveys of land cover types and activities of humans, livestock and wildlife (Section 2.3) that were used for the analyses described in sections 2.4, 2.5 and 2.7 https://doi.org/10.5281/zenodo.11099625

**Supplementary File 9**; Figures S9.1 to S9.10: Maps of the ILUMA WMA, respectively illustrating the geographic distribution of all the various human activities surveyed as described in section 2.3 https://doi.org/10.5281/zenodo.11099630

**Supplementary File 10;** Tables S10.1 to S10.3 detailing the results of univariate and multivariate generalized linear modelling analyses of scaled human activity indicators as respective determinants of the estimated Species Richness Index (SRI), Simpson’s Index of Diversity (SID) and Objective Wild Animal Community Integrity Index (OWACII) for each camp surveyed. Integrity https://doi.org/10.5281/zenodo.11099636

**Supplementary File 11;** Supplementary figures S11.1 to S11.5; Maps illustrating how the distributions of the values for all the significant independent variables that were included in the final multivariate model detailed in table 1, except for the extent of human settlement (See figure 8), compared with those for the Subjective Natural Ecosystem Integrity Index (SNEII) scores that were used as the dependent variable in that model https://doi.org/10.5281/zenodo.11099639

**Supplementary File 12;** Supplementary figures S12.1 to S12.5; Dot plots and violin plots respectively illustrating the absolute frequency and probability density distributions of all the various recorded signs of activity by wild animals, livestock, work animals and humans during the surveys described in section 2.3 https://doi.org/10.5281/zenodo.11099645

