## Supplementary File 1 for "An effective model for community-based conservation around authorized fishing settlements inside a devolved Wildlife Management Area in southern Tanzania" (from another version of this preprint: version 1, 2024-07-31)

**Table 1.** The number, name, location, coordinates, and habitat type of each camp is described. Also included here is the quadrant assigned to the camp and the number of times it was surveyed.

| **Camp Number** | **Camp Name** | **Camp Location and Coordinates** | **No. of Times Surveyed** | **Quadrant** | **Habitat Type** |
| --- | --- | --- | --- | --- | --- |
| 1 | Msakamba | -8.25981S, 36.85936E  ILUMA WMA | 3 | Southeast | Recovering miombo woodland on a large river. |
| 2 | Msiba wa Deo | -8.25821S, 36.88524E  ILUMA WMA | 3 | Southeast | Intact miombo woodland on a large river. |
| 3 | Bwawa la Nyati | -8.29598S, 36.88732E  ILUMA WMA | 3 | Southeast | Intact miombo woodland surrounding a large waterhole. |
| 4 | Bwawa la Nandete | -8.3467S, 36.9024E  ILUMA WMA | 3 | Southeast | Degraded miombo woodland surrounding a large waterhole. |
| 5 | Korongo la Bundu | -8.37768S, 36.90163E  ILUMA WMA | 3 | Southeast | Degraded miombo woodland on a large river. |
| 6 | Bwawa la Namamba | -8.34931S, 36.81653E  ILUMA WMA | 3 | Southeast | Degraded miombo woodland surrounding a large waterhole. |
| 7 | Bwawa la Chakacheni | -8.31649S, 36.81507E  ILUMA WMA | 3 | Southeast | Degraded miombo woodland surrounding a large waterhole. |
| 8 | Bwawa la Njuju | -8.24204S, 36.86268E  ILUMA WMA | 3 | Northeast | Intact miombo woodland surrounding a large waterhole. |
| 9 | Bwawa la Chamvi | -8.22176S, 36.89241E  ILUMA WMA | 2 | Northeast | Intact miombo woodland surrounding a large waterhole. |
| 10 | Bwawa la Miembeni | -8.22401S, 36.87518E  ILUMA WMA | 1 | Northeast | Mostly intact miombo woodland surrounding a waterhole. |
| 11 | Kisima cha Seba | -8.20749S, 36.85738E  ILUMA WMA | 2 | Northeast | Intact miombo woodland adjacent to a small waterhole. |
| 12 | Bwawa la Maya | -8.19492S, 36.88045E  ILUMA WMA | 3 | Northeast | Transition zone between miombo woodland and groundwater forest surrounding a large waterhole. |
| 13 | Bwawa la Mrope | -8.16321S, 36.88341E  ILUMA WMA | 2 | Northeast | Dense groundwater forest surrounding a large waterhole. |
| 14 | Mikeregembe | -8.15048S, 36.87539E  ILUMA WMA | 3 | Northeast | Human settlement on the banks of the Kilombero river, open but bordering groundwater forest. |
| 15 | Mdalangwila | -8.16041S, 36.83887E  ILUMA WMA | 3 | Northeast | Human settlement on the banks of the Kilombero river, open but bordering groundwater forest. |
| 16 | Bwawa la Muamachi | -8.38785S, 36.74629E  Outside ILUMA | 3 | Southwest | Human settlement surrounding a large waterhole. |
| 17 | Tuliza Moyo | -8.35954S, 36.74821E  Outside ILUMA | 3 | Southwest | Human settlement along a large river. |
| 18 | Mavimba Porini | -8.33982S, 36.74628E  Outside ILUMA | 3 | Southwest | Human settlement expanding along a large river and surrounding a large waterhole. |
| 19 | Bwawa la Selesussi | -8.33064S, 36.76951E  ILUMA WMA | 3 | Southwest | Degraded miombo woodland surrounding a large waterhole. |
| 20 | Makingi | -8.27593S, 36.74737E  Outside ILUMA | 3 | Southwest | Human settlement along a large river. |
| 21 | Bwawa la Mpunga | -8.27426S, 36.81946E  ILUMA WMA | 3 | Southwest | Highly degraded miombo woodland surrounding a large waterhole. |
| 22 | Kisaki | -8.24595S, 36.7669E  Outside ILUMA | 2 | Northwest | Largest human settlement, at the base of a large hill adjacent to a spring and large waterhole. |
| 23 | Uwanja wa Ndege | -8.23448S, 36.80218E  Outside ILUMA | 2 | Northwest | Human settlement along a large river. |
| 24 | Bwawa la Mkwajuni | -8.21681S, 36.81141E  ILUMA WMA | 2 | Northwest | Intact miombo woodland surrounding a degraded large waterhole. |
| 25 | Bwawa la Mamba Luhogi | -8.19352S, 36.78527E  ILUMA WMA | 2 | Northwest | Intact miombo woodland surrounding a highly degraded large waterhole. |
| 26 | Funga | -8.16924S, 36.776E  ILUMA WMA | 2 | Northwest | Human settlement on the banks of the Kilombero river, open but bordering intact miombo woodland. |
| 27 | Bwawa la Mlenda | -8.20037S, 36.82132E  ILUMA WMA | 2 | Northwest | Intact miombo woodland surrounding a large waterhole. |
| 28 | Bwawa la Semka | -8.20436S, 36.84023E  ILUMA WMA | 2 | Northwest | Largely intact groundwater forest surrounding a large waterhole. |
| 29 | Bwawa la Simba | -8.31298S, 36.94381E  Nyerere NP | 1 | Nyerere | Transition zone of miombo woodland and acacia savanna surrounding a waterhole. |
| 30 | Kiboko Zanzibar | -8.26313S, 37.00293E  Nyerere NP | 1 | Nyerere | Open acacia savanna on the banks of the Kilombero river. |
| 31 | Zanzibar | -8.2565S, 36.98337E  Nyerere NP | 1 | Nyerere | Transition zone of miombo woodland and acacia savanna on the banks of the Kilombero river. |
| 32 | Bwawa la Moto | -8.2741S, 36.93866E  Nyerere NP | 1 | Nyerere | Transition zone of miombo woodland and acacia savanna surrounding a waterhole. |
