## Supplementary File 2 for "An effective model for community-based conservation around authorized fishing settlements inside a devolved Wildlife Management Area in southern Tanzania" (from another version of this preprint: version 1, 2024-07-31)

**Table 2.** Results of various ecological indices calculated for each location (camp). Cluster Analysis of all activities, entities, and land cover attributes. Species Richness of all recorded wildlife data. Simpson’s Index of Diversity of all recorded wildlife data. Subjective, opinion-based Ecosystem Integrity Score. Natural Community Integrity Index based on PCA of all recorded parameters for all wildlife data. Natural Ecosystem Integrity Index PCA based index of all recorded parameters of humans, livestock, and wildlife.

| Camp | Cluster^a^ | Location | Historic Natural Land Cover | Species Richness Index^b^ | | Simpson’s Index of Diversity^c^ | | Subjective Natural Ecosystem Integrity Index^d^ | | Objective Wild Animal Community Index^e^ | | Objective Natural Ecosystem Integrity Index^f^ | |
| --- | --- | --- | --- | --- | --- | --- | --- | --- | --- | --- | --- | --- | --- |
|  |  |  |  | Value | Rank^g^ | Value | Rank^g^ | Value | Rank^g^ | Value | Rank^g^ | Value | Rank^g^ |
| 1: Msakamba | Intact | Inside ILUMA WMA | Miombo Woodland | 16 | 6 | 0.952 | 4 | 70 | 17 | -0.507 | 20 | -0.390 | 14 |
| 2: Msiba wa Deo | Intact | Inside ILUMA WMA | Miombo Woodland | 20 | 2 | 0.934 | 12 | 84 | 9 | 0.338 | 6 | -0.478 | 6 |
| 3: Bwawa la Nyati | Intact | Inside ILUMA WMA | Miombo Woodland | 17 | 5 | 0.957 | 1 | 85 | 8 | 0.250 | 8 | -0.439 | 9 |
| 4: Bwawa la Nandete | Disturbed | Inside ILUMA WMA | Miombo Woodland | 16 | 6 | 0.956 | 2 | 35 | 23 | -0.349 | 15 | -0.370 | 17 |
| 5: Korongo la Bundu | Disturbed | Inside ILUMA WMA | Miombo Woodland | 18 | 4 | 0.951 | 5 | 30 | 24 | -0.468 | 18 | -0.352 | 20 |
| 6: Bwawa la Namamba | Disturbed | Inside ILUMA WMA | Miombo Woodland | 12 | 10 | 0.889 | 18 | 40 | 22 | -0.619 | 28 | -0.347 | 21 |
| 7: Bwawa la Chakacheni | Disturbed | Inside ILUMA WMA | Miombo Woodland | 6 | 13 | 0.652 | 26 | 15 | 26 | -0.594 | 24 | -0.333 | 25 |
| 8: Bwawa la Njuju | Intact | Inside ILUMA WMA | Miombo Woodland | 19 | 3 | 0.952 | 3 | 81 | 12 | -0.169 | 12 | -0.411 | 11 |
| 9: Bwawa la Chamvi | Intact | Inside ILUMA WMA | Miombo Woodland | 19 | 3 | 0.950 | 6 | 86 | 7 | -0.004 | 10 | -0.418 | 10 |
| 10: Bwawa la Miembeni | Intact | Inside ILUMA WMA | Miombo Woodland | 15 | 7 | 0.933 | 13 | 68 | 18 | 0.035 | 9 | -0.443 | 8 |
| 11: Kisima cha Seba | Intact | Inside ILUMA WMA | Miombo Woodland | 9 | 12 | 0.882 | 19 | 82 | 11 | -0.410 | 17 | -0.369 | 18 |
| 12: Bwawa la Maya | Intact | Inside ILUMA WMA | Miombo Woodland | 22 | 1 | 0.928 | 14 | 87 | 6 | 0.296 | 7 | -0.445 | 7 |
| 13: Bwawa la Mrope | Intact | Inside ILUMA WMA | Groundwater Forest | 12 | 10 | 0.893 | 17 | 90 | 5 | -0.244 | 13 | -0.383 | 15 |
| 14: Mikeregembe | Intact | Inside ILUMA WMA | Groundwater Forest | 17 | 5 | 0.947 | 9 | 75 | 15 | -0.500 | 19 | -0.382 | 16 |
| 15: Mdalangwila | Disturbed | Inside ILUMA WMA | Groundwater Forest | 22 | 1 | 0.941 | 10 | 72 | 16 | -0.260 | 14 | -0.399 | 13 |
| 16: Bwawa la Muamachi | Disturbed | Outside ILUMA WMA | Miombo Woodland | 4 | 15 | 0.574 | 27 | 6 | 28 | -0.603 | 26 | 1.325 | 30 |
| 17: Tuliza Moyo | Disturbed | Outside ILUMA WMA | Miombo Woodland | 5 | 14 | 0.340 | 30 | 4 | 30 | -0.581 | 23 | 1.082 | 28 |
| 18: Mavimba Porini | Disturbed | Outside ILUMA WMA | Miombo Woodland | 4 | 15 | 0.159 | 32 | 10 | 27 | -0.568 | 22 | 0.065 | 26 |
| 19: Bwawa la Selessusi | Disturbed | Outside ILUMA WMA | Miombo Woodland | 11 | 11 | 0.740 | 25 | 25 | 25 | -0.749 | 31 | -0.337 | 24 |
| 20: Makingi | Disturbed | Outside ILUMA WMA | Miombo Woodland | 2 | 16 | 0.234 | 31 | 0 | 32 | -0.597 | 25 | 1.185 | 29 |
| 21: Bwawa la Mpunga | Disturbed | Inside ILUMA WMA | Miombo Woodland | 12 | 10 | 0.785 | 23 | 55 | 19 | -0.721 | 30 | -0.344 | 22 |
| 22: Kisaki | Disturbed | Outside ILUMA WMA | Miombo Woodland | 5 | 14 | 0.411 | 29 | 5 | 29 | -0.607 | 27 | 3.805 | 32 |
| 23: Uwanja wa Ndege | Disturbed | Outside ILUMA WMA | Miombo Woodland | 6 | 13 | 0.504 | 28 | 3 | 31 | -0.621 | 29 | 2.781 | 31 |
| 24: Bwawa la Mkwajuni | Disturbed | Inside ILUMA WMA | Miombo Woodland | 11 | 11 | 0.841 | 21 | 45 | 21 | -0.529 | 21 | -0.340 | 23 |
| 25: Bwawa la Mamba Luhogi | Disturbed | Inside ILUMA WMA | Miombo Woodland | 15 | 7 | 0.836 | 22 | 48 | 20 | -0.850 | 32 | 0.144 | 27 |
| 26: Funga | Intact | Inside ILUMA WMA | Groundwater forest | 13 | 9 | 0.754 | 24 | 77 | 14 | -0.401 | 16 | -0.366 | 19 |
| 27: Bwawa la Mlenda | Disturbed | Inside ILUMA WMA | Miombo Woodland | 15 | 7 | 0.879 | 20 | 83 | 10 | -0.162 | 11 | -0.408 | 12 |
| 28: Bwawa la Semka | Intact | Inside ILUMA WMA | Miombo Woodland | 16 | 6 | 0.928 | 15 | 80 | 13 | 0.587 | 5 | -0.488 | 5 |
| 29: Bwawa la Simba | Intact | Nyerere National Park | Groundwater Forest | 11 | 11 | 0.910 | 16 | 97 | 3 | 2.356 | 3 | -0.549 | 2 |
| 30: Kiboko Zanzibar | Intact | Nyerere National Park | Acacia Savanna | 16 | 6 | 0.935 | 11 | 100 | 1 | 2.920 | 1 | -0.519 | 3 |
| 31: Zanzibar | Intact | Nyerere National Park | Miombo Woodland | 14 | 8 | 0.949 | 7 | 92 | 4 | 1.606 | 4 | -0.489 | 4 |
| 32: Bwawa la Moto | Intact | Nyerere National Park | Miombo Woodland | 18 | 4 | 0.948 | 8 | 99 | 2 | 2.724 | 2 | -0.590 | 1 |

^a^ Interpreted results of a cluster analysis performed on every variable collected, including all human, livestock, and wildlife activity, as well as conditions such as ground type, vertical grass height and horizontal density of tree, bush and shrub cover.

^b^ Defined as *S* the sum of the total number of species recorded in a particular area (each camp).

^c^ Calculates the probability of selecting two individuals from different species within a defined area (camp) and calculated as SDI=1- D, where D = $\sum iPi$ ^2^, where the species proportion(the proportion of a particular species divided by the total number of all species observations) (*P_i_*) is calculated as *P_i_* = *x_i_*/*n*, where *x_i_* is the total number of individuals of species *i* recorded and *n* is the total number of individuals from all species recorded.

^d^ Based on personal, subjective opinions of the three investigators based on their impressions over the time spent in the field, with each camp being given a consensus score from 0 to 100%, with 100% representing completely pristine conditions with no evidence of human activities and 0% representing highly degraded areas that have been completely converted into fully domesticated habitats.

^e^ Scaled results of Principal Component Analysis of all parameters recorded for wild herbivores, rodents, carnivores, primates, and prosimians.

^f^ Scaled results of Principal Component Analysis of all parameters recorded for human, livestock, and wild animals.

^g^ Locations ranked from the most species rich, biodiverse, or ecologically intact to the least, based on values from each index.
