## Supplementary File 3 for "An effective model for community-based conservation around authorized fishing settlements inside a devolved Wildlife Management Area in southern Tanzania" (from another version of this preprint: version 1, 2024-07-31)

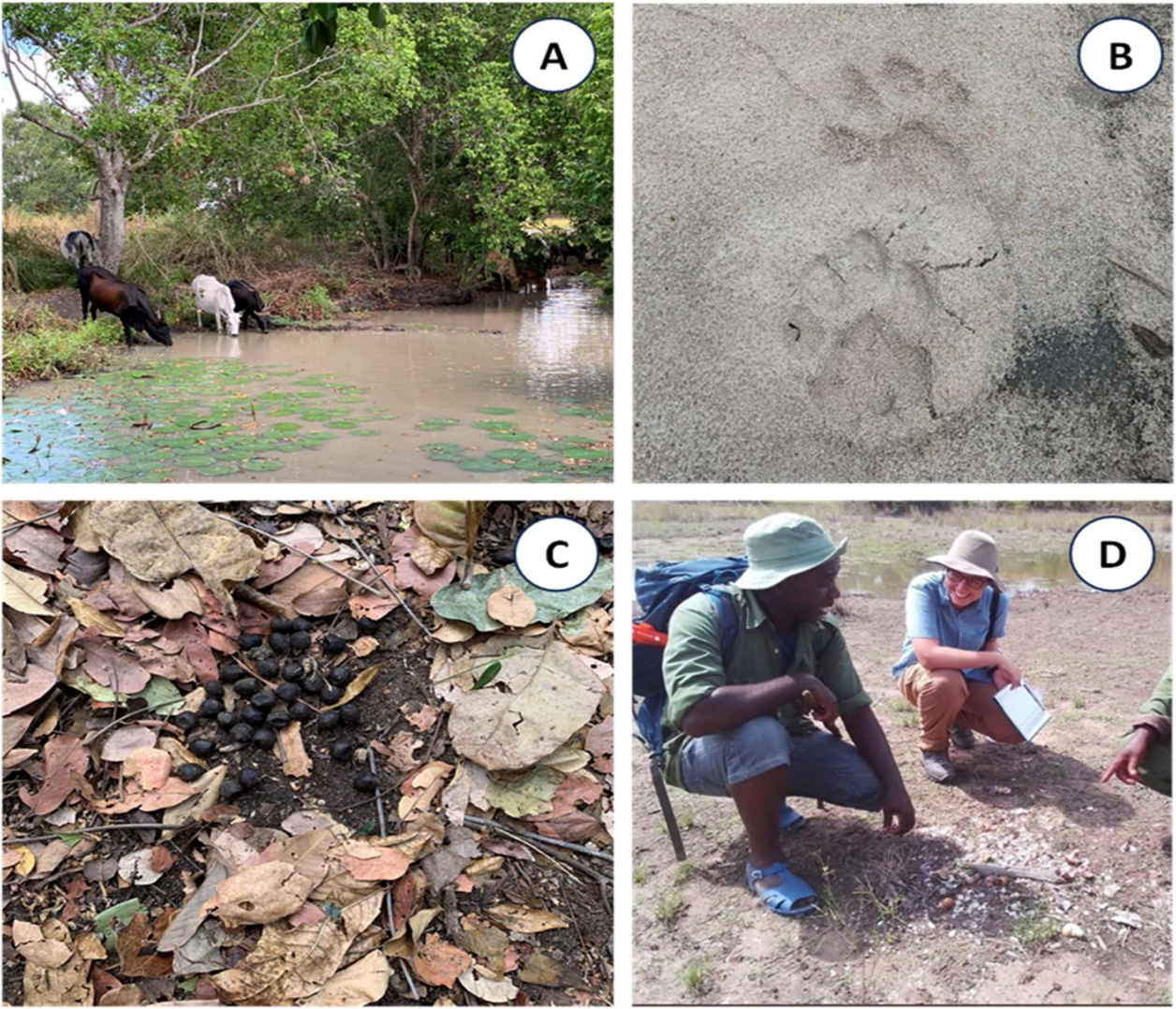

**Figure 10.** Images illustrating different ways in which livestock and/or human recordings (direct sighting, tracks, spoor or other signs) were detected. Panel A shows a direct sighting of cattle. Panel B shows an observation rarely detected, namely fresh lion prints on damp sand which resulted in perfectly clear tracks and an unambiguous identification of the animal and age of the tracks. Contrasting this, panel C shows one of the most frequently recorded detections throughout the entire survey - Hartebeest spoor, 3 days old. Panel D displays an example of “other signs”, here broken shells and a stick used as an anvil by water mongoose to obtain snails and shellfish is shown. This sign was frequently recorded throughout the study.
