## Supplementary File 4 for "An effective model for community-based conservation around authorized fishing settlements inside a devolved Wildlife Management Area in southern Tanzania" (from another version of this preprint: version 1, 2024-07-31)

**Supp 4.** Complete list of all wild herbivores, wild carnivores, wild primates and prosimians and wild rodents sampled in this study.

| **Wild Herbivores.** | **Wild Carnivores.** | **Wild Primates and Prosimians.** |
| --- | --- | --- |
| Bohor Reedbuck (*Redunca redunca wardi*) | Lion (*Panthera leo*) | Lesser Bush Baby (*Galago senegalensis*) |
| Common Waterbuck (*Kobus ellipsiprymnus ellipsiprymnus*) | Leopard (*Panthera pardus*) | Greater Bush Baby (*Galago crassicaudatus*) |
| Puku (*Kobus vardoni*) | African Wild Cat (*Felis libyca*) | Yellow Baboon (*Papio cynocephalus cynocephalus*) |
| Bushbuck (*Tragelaphus scriptus*) | African Civet (*Civettictis civetta*) | Vervet Monkey (*Cercopithecus aethiops*) |
| Common Eland (*Taurotragus oryx*) | Genet (*Genetta spp.*) | Blue Monkey (*Cercopithecus mitis*) |
| Greater Kudu (*Tragelaphus strepsiceros*) | Spotted Hyena (*Crocuta Crocuta*) |  |
| Hartebeest (*Alcelaphus buselaphus*) | African Wild Dog (*Lycaon pictus*) | **Wild Rodents.** |
| Wildebeest (*Coonochaetes taurinus*) | Side-Striped Jackal (*Canis* *adustus*) | Porcupine (*Hystrix cristata*) |
| Sable (*Hippotragus niger*) | Slender Mongoose (*Herpestes sanguineus*) | Giant Cane Rat (*Thryonomys swinderainus*) |
| Warthog (*Phacochoerus aethiopicus*) | Banded Mongoose (*Mungos mungo*) |  |
| Bushpig (*Potamochoerus porcus*) | Water Mongoose (*Atilax paludinosus*) | **Macroscelidea.** |
| Hippopotamus (*Hippopotamus amphibius*) | White-Tailed Mongoose (*Ichneumia albicauda*) | Sengi (*Petrodromus tetradactylus*) |
| Elephant (*Loxodonta Africana*) | Honey Badger (*Mellivora capensis*) |  |
| African Buffalo (*Syncerus caffer*) | African Clawless Otter (*Aonyx capensis*) | **Tubulidentata.** |
| Plains Zebra (*Equus burchelli*) |  | Aardvark (*Orycteropus afer*) |
| Suni (*Neotragus moschatus*) |  |  |
| Common Duiker (*Sylvicapra grimmia*) |  |  |
| Natal Red Duiker (*Cephalophus natalensis*) |  |  |
| Impala (*Aepyceros melampus*) |  |  |
| Ugogo Dikdik (*Madoqua kirkii thomasi*) |  |  |
| Sharpe’s Grysbok (*Raphicerus sharpei*) |  |  |
