## Supplementary File 5 for "An effective model for community-based conservation around authorized fishing settlements inside a devolved Wildlife Management Area in southern Tanzania" (from another version of this preprint: version 1, 2024-07-31)

| **Entity or Activity Observed (EAO)** | **Observation Category (OC)** | **Additional Observation Attributes (AOA)** | **Observation Count or Class (OCC)** |
| --- | --- | --- | --- |
| **1X** “As you go” instances of specific human land and/or natural resources usages. |  |  |  |
| **11**: Livestock Herding |  |  | **1**: Active, **2**: Recent, **3**: Old, **9**: Not determined or recorded |
| **12**: Charcoal Burning |  |  | **1**: Active, **2**: Recent, **3**: Old, **9**: Not determined or recorded |
| **13**: Timber Harvesting |  |  | **1**: Active, **2**: Recent, **3**: Old, **9**: Not determined or recorded |
| **14**: Fishing |  |  | **1**: Active, **2**: Recent, **3**: Old, **9**: Not determined or recorded |
| **15**: Hunting |  |  | **1**: Active, **2**: Recent, 3: Old, **9**: Not determined or recorded |
| **16**: Human Settlement |  | **1,2,3…10** structures [n], 11-20 structures **[20],** 21-50 structures **[50],** >50 structures **[100]**, not determined or recorded **[9]** | **1**: Active, **2**: Recent, **3**: Old, **9**: Not determined or recorded |
| **17**: Water Body |  |  | **1**: Bwawa. **2**: (Ma)dimbwi outside korongo **3:** (Ma)dimbwi inside korongo **4**: Flowing korongo **5**: Bonde |
| **18:** Human dug well |  |  | **1**: Active, **2**: Recent, **3**: Old, **9**: Not determined or recorded |
| **19:** Meat Poaching |  |  | **1:** Active**, 2:** Recent, **3:** Old**, 9:** Not determined or recorded |
| **2X**: “As you go” sightings, tracks and signs of humans and animals |  |  |  |
| **21**: Humans | 1: Direct sighting, 2: Tracks, 3: Spoor, **4**: Other signs (Explain in notes) | **1**: ILUMA personnel, **2**: NNP Rangers, **3**: Hunting company staff, **4**: Other authorized personnel from ILUMA and other authorized institutions, **5**: Community legal presence inside conservation area, **6**: Community legal presence outside the conservation area, **7**: Charcoal transport, **8**: Timber harvesting, **9**: Timber transport, **10**: Fish harvesting, **11**: Fish transport, **12**: Community illegal presence inside the conservation area, **13:** Charcoal burner. | Distance (m) from track for direct observations, or age for tracks and signs: **1**: Active, **2**: Recent, **3**: Old, **9**: Not determined or recorded |
| **22**: Livestock and work animals | **1**: Direct sighting, **2**: Tracks, **3**: Spoor, **4**: Other | **1**: Cattle, **2**: Goats, **3**: Sheep, **4**: Rooster, **5**: Chicken, **6**: Dog | Distance (m) from track for direct observations, or age for tracks and signs: **1**: Active, **2**: Recent, **3**: Old, **9**: Not determined or recorded |
| **23**: Wild Herbivores | **1**: Direct sighting, **2**: Tracks, **3**: Spoor, **4**: Other signs (Explain in notes) | 1X: Reduncini (**11**: Reedbuck, **12**: Waterbuck, **13**: Puku), 2X: Spiral Horned Antelopes (**21**: Bushbuk, **22**: Eland, **23**: Greater Kudu) 3X: Alcephalines (**31**: Hartebeest, **32**: Wildebeest), 4X: Hippotragini (**41**: Sable Antelope), 5X: Suidae (**51**: Warthog, **52**: Bushpig), 6X Plantigrade non-ungulates (**61**: Hippo, **62**: Elephant), 7X: Bovinae (**71**: African Buffalo), 8X: Equidae (**81**: Plains Zebra), 9X: Neotragini (**91**: Suni), 10X: Cephalophini (**101**: Common Duiker, **102**: Red Duiker), 11X: Aepycerotini (**111**: Common Impala), 12X: Madoquini (**121**: Dikdik), 13X: Raphicerini (**131**: Sharpe's Grysbok) | Distance (m) from track for direct observations, or age for tracks and signs: **1**: Active, **2**: Recent, **3**: Old, **9**: Not determined or recorded |
| **24**: Wild Carnivores | 1: Direct sighting, 2: Tracks, 3: Spoor, 4: Other signs (Explain in notes) | 1X: Pantherinae (**11**: Lion, **12**: Leopard), 2X: Felidae (**21**: African wild cat), 3X: Viverridae (**31**: African Civet, **32**: Cape Genet), 4X: Hyaenidae (**41**: Spotted Hyena), 5X: Canidae (**51**: African wild dog, **52**: Side - Striped Jackal), 6X: Herpestidae (**61**: Slender Mongoose, **62**: Banded Mongoose **63:** Water Mongoose, **64:** White Tailed Mongoose), 7X: Mustelids (**71**: Honey Badger), 8X Lutrinae ( **81**: African Clawless Otter) | Distance (m) from track for direct observations, or age for tracks and signs: **1**: Active, **2**: Recent, **3**: Old, **9**: Not determined or recorded |
| **25**: Wild Primates and Prosimians | **1**: Direct sighting, **2**: Tracks, **3**: Spoor, **4**: Other | **1**: Lesser Galago, **2**: Greater Galago, **3**: Baboon, **4**: Vervet, **5**: Blue Monkey | Distance (m) from track for direct observations, or age for tracks and signs: **1**: Active, **2**: Recent, **3**: Old, **9**: Not determined or recorded |
| **26**: Rodents | **1**: Direct sighting, **2**: Tracks, **3**: Spoor, **4**: Other | 1: Hystricidae (**11**: Porcupine, **12**: Brush tailed porcupine) | Distance (m) from track for direct observations, or age for tracks and signs: **1**: Active, **2**: Recent, **3**: Old, **9**: Not determined or recorded |
| **27:** Macroscelidea | **1**: Direct sighting, **2**: Tracks, **3**: Spoor, **4**: Other | 1: Rhynchoninae (**11:** Sengei) | Distance (m) from track for direct observations, or age for tracks and signs: **1**: Active, **2**: Recent, **3**: Old, **9**: Not determined or recorded |
| **28:** Tubulidentata | **1**: Direct sighting, **2**: Tracks, **3**: Spoor, **4**: Other | 1: Orycteropodidae (**11:** Aardvark) | Distance (m) from track for direct observations, or age for tracks and signs: **1**: Active, **2**: Recent, **3**: Old, **9**: Not determined or recorded |
| **3X**: Summary impressions of land cover attributes averaged over entire segment |  |  |  |
| **31**: Rice farming |  |  | **1**: 0%, **2**: 1 - 25%, **3**: 26 - 50%, **4**: 51 - 75%, **5**: 76 - 100% |
| **32**: Other Tillage Crops |  |  | **1**: 0%, **2**: 1 - 25%, **3**: 26 - 50%, **4**: 51 - 75%, **5**: 76 - 100% |
| **33**: Grass and forb height |  |  | **1:** <1ft, **2:** 1 - 3ft, **3:** 3 -6ft, **4:** 6ft+ |
| **34:** Visibility of obstruction by Shrubs, Bushes and Trees |  |  | **1**: 0%, **2**: 1 - 25%, **3**: 26 - 50%, **4**: 51 - 75%, **5**: 76 - 100% |
| **35:** Ground Type |  |  | **1**: Hard & Dry, **2:** Sand**, 3:** Damp Soil, **4**: Mud, **5:** Water**, 6:** Rock**, 7:** Gravel & Pebbles |
