## Supplementary File 6 for "An effective model for community-based conservation around authorized fishing settlements inside a devolved Wildlife Management Area in southern Tanzania" (from another version of this preprint: version 1, 2024-07-31)

**Supp 6.** Defining terms used in Data Dictionary.

**Direct sighting:** Live animal or human actually seen by one or more members of the field team.

**Tracks:** Prints (animal or human) seen by the data recorder on the ground on the immediate path being followed.

**Spoor:** Faecal excrement seen by the data recorder on the on the immediate path being followed.

**Other signs:** Any sign of animal activity that undoubtedly links to a specific species (elephant pulling bark from a tree or water mongoose cracking shells for example).

**Bwawa:** Stagnant water body, may be permanent or ephemeral.

**(Ma)dimbwi:** Puddle(s).

**Korongo:** River or riverbed.

**Bonde:** Flooded valley.
