## Supplementary File 7 for "An effective model for community-based conservation around authorized fishing settlements inside a devolved Wildlife Management Area in southern Tanzania" (from another version of this preprint: version 1, 2024-07-31)

| **Form Type (FT):** Land Use Observations (LUO) | | | | **Date (DT):** / / | | **GPS Identifier (GID):** | |  | **Form Serial Number (SEN):** |
| --- | --- | --- | --- | --- | --- | --- | --- | --- | --- |
| **Transect Start Camp Number (TSCNO):** | | | | **Transect Start Camp Name or Description (TSCND):** | | | | |  |
| **Transect Segment Number (TSN):** | | |  |  | **Transect Start GPS Way Point Number (TSGWPN):** | | | | |
| **Transect Segment Page Number (TSPN):** / | | | |  | **Transect End GPS Way Point Number (TEGWPN):** | | | | |
| **Form Row (FR)** | **Entity or Activity Observed (EAO)** | **Observation Category (OC)** | **Additional observation attributes (AOA)** | **Observation count or class (OCC)** | **Observation Notes (ONO):** Specify details that add qualitative insight or enable post-hoc classification | | | | |
| 1 |  |  |  |  |  |  |  |  |  |
| 2 |  |  |  |  |  |  |  |  |  |
| 3 |  |  |  |  |  |  |  |  |  |
| 4 |  |  |  |  |  |  |  |  |  |
| 5 |  |  |  |  |  |  |  |  |  |
| 6 |  |  |  |  |  |  |  |  |  |
| 7 |  |  |  |  |  |  |  |  |  |
| 8 |  |  |  |  |  |  |  |  |  |
| 9 |  |  |  |  |  |  |  |  |  |
| 10 |  |  |  |  |  |  |  |  |  |
| 11 |  |  |  |  |  |  |  |  |  |
| 12 |  |  |  |  |  |  |  |  |  |
| 13 |  |  |  |  |  |  |  |  |  |
| 14 |  |  |  |  |  |  |  |  |  |
| 15 |  |  |  |  |  |  |  |  |  |
| 16 |  |  |  |  |  |  |  |  |  |
| 17 |  |  |  |  |  |  |  |  |  |
| 18 |  |  |  |  |  |  |  |  |  |
| 19 |  |  |  |  |  |  |  |  |  |
| 20 |  |  |  |  |  |  |  |  |  |
| 21 |  |  |  |  |  |  |  |  |  |
| 22 |  |  |  |  |  |  |  |  |  |
| 23 |  |  |  |  |  |  |  |  |  |
| 24 |  |  |  |  |  |  |  |  |  |
| 25 |  |  |  |  |  |  |  |  |  |

**Supp 7.** Data collection sheet developed during preliminary investigation in October and November 2021 and used for all data collection in both transect segment surveys and camp radial surveys throughout the course of the study.
