## Supplementary figures and images for "An effective model for community-based conservation around authorized fishing settlements inside a devolved Wildlife Management Area in southern Tanzania" (from another version of this preprint: version 1, 2024-07-31)

### Supplementary File 9

**Supp 9**. Maps of ILUMA WMA displaying human activities recorded.
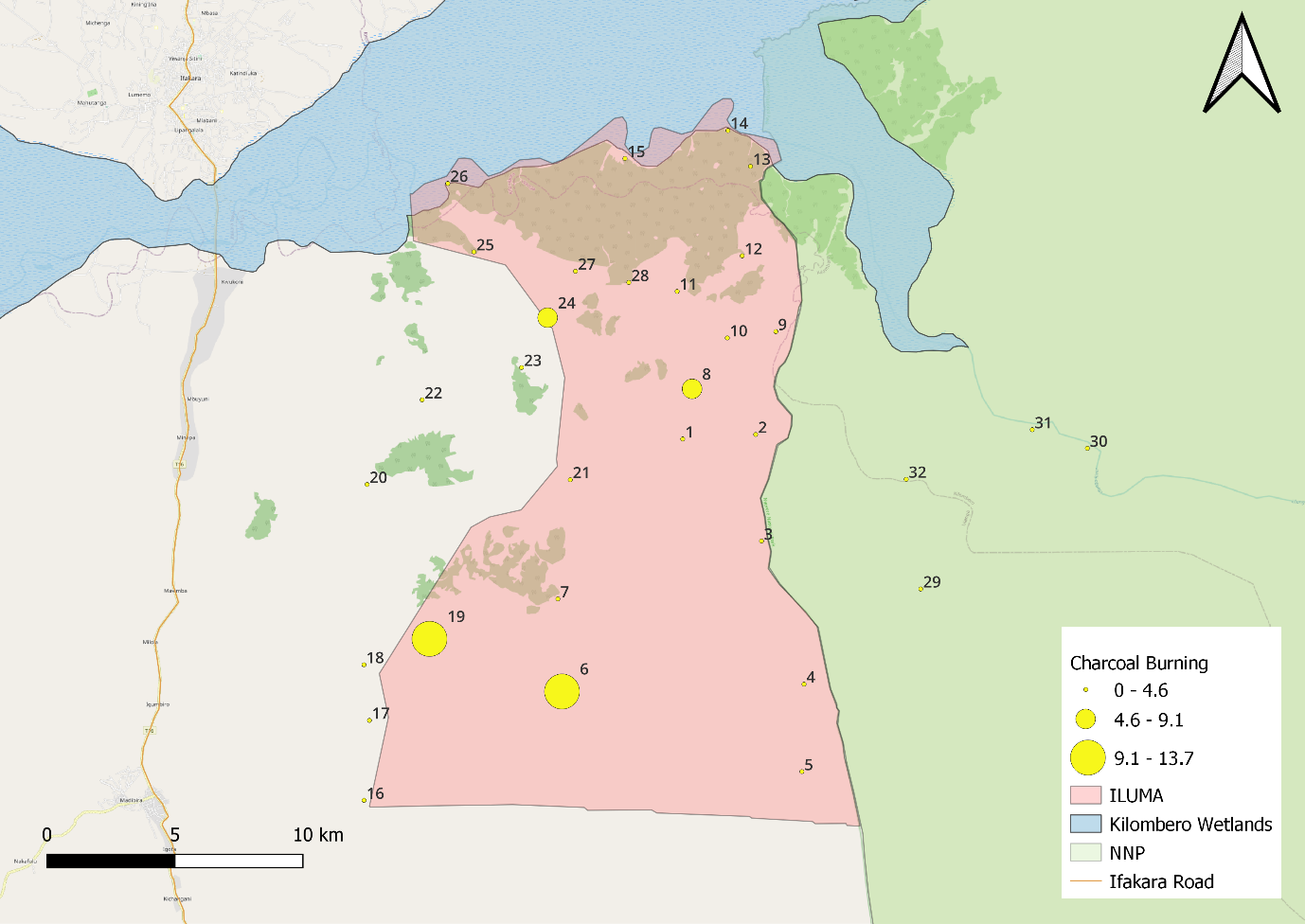

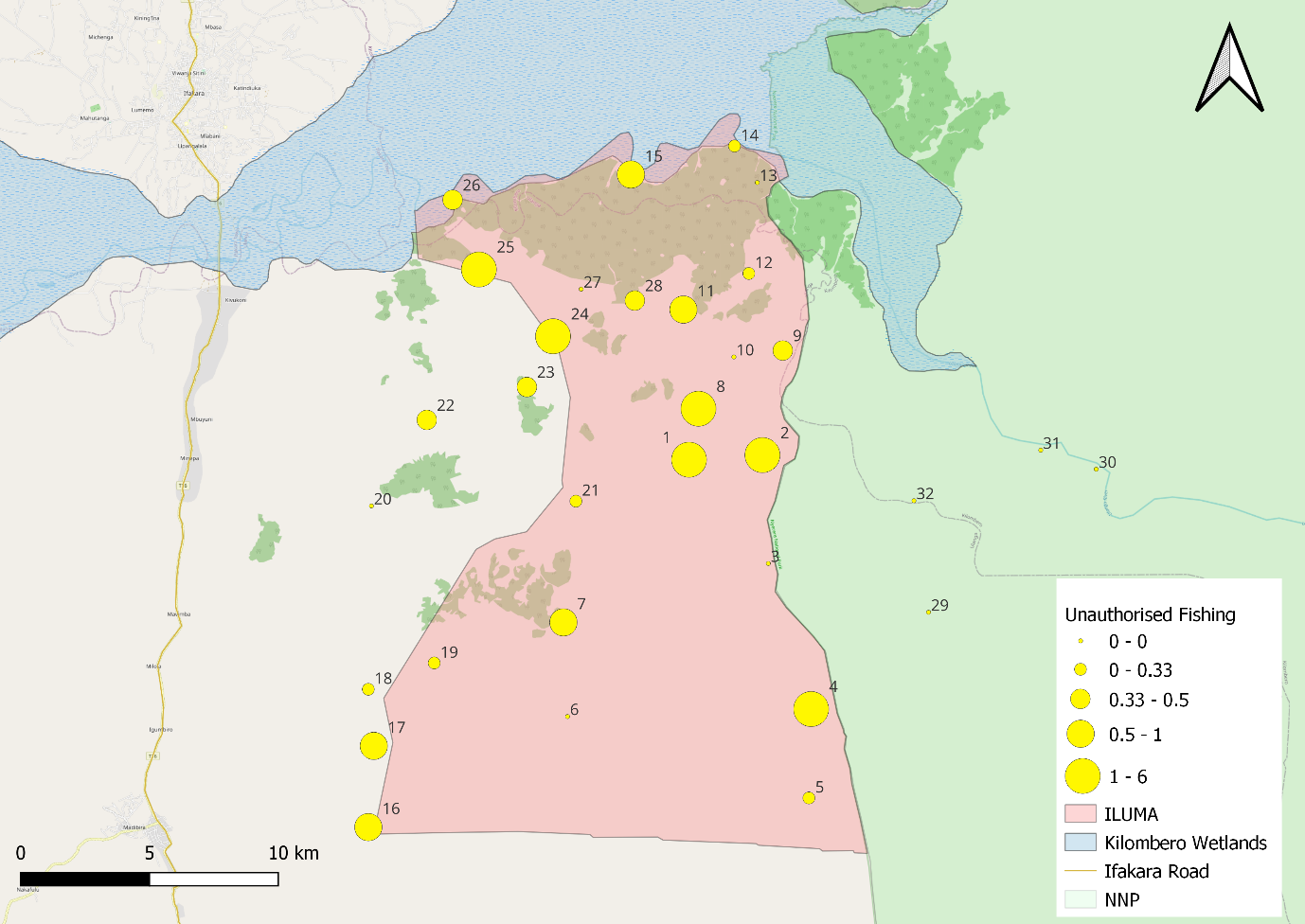

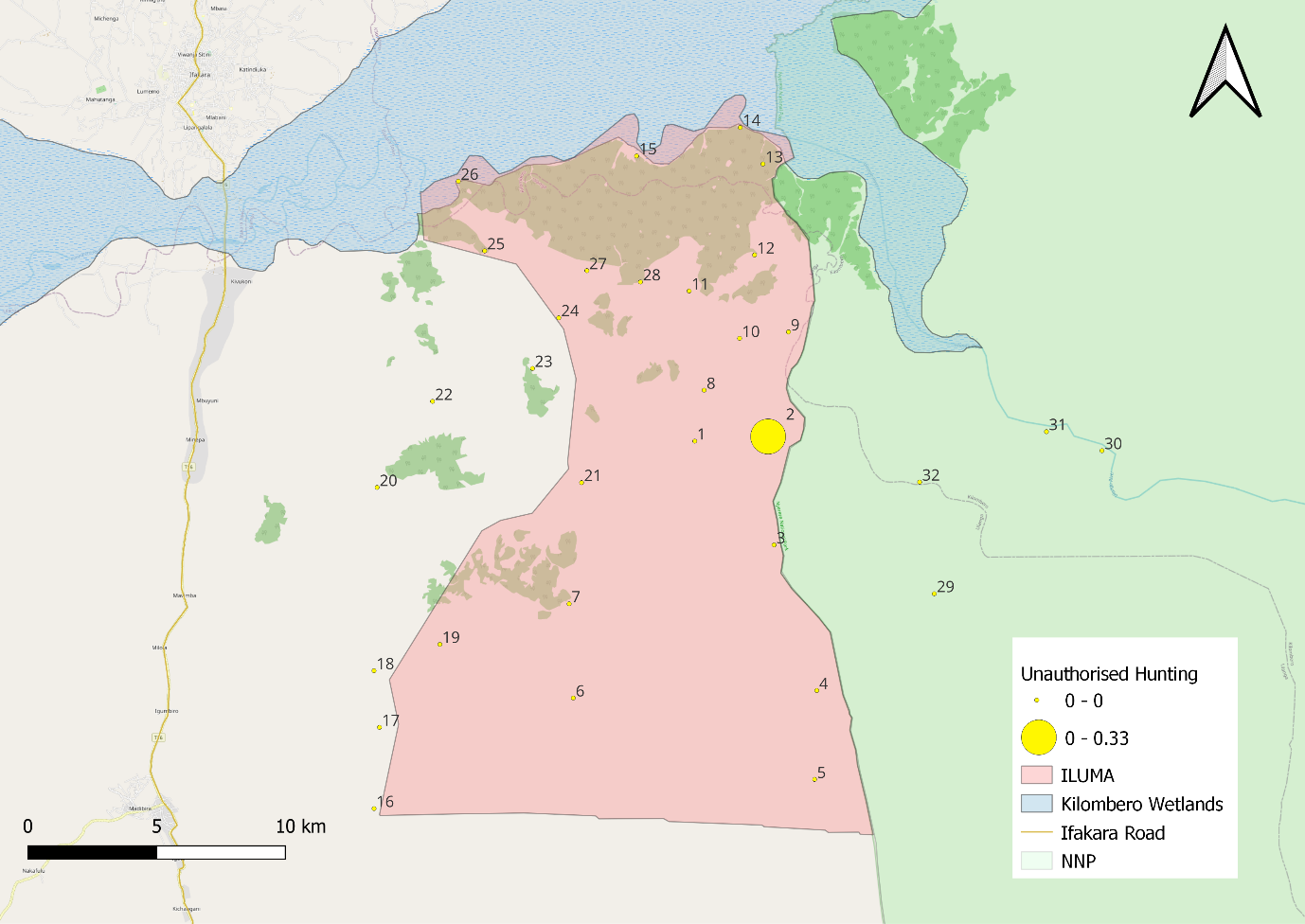

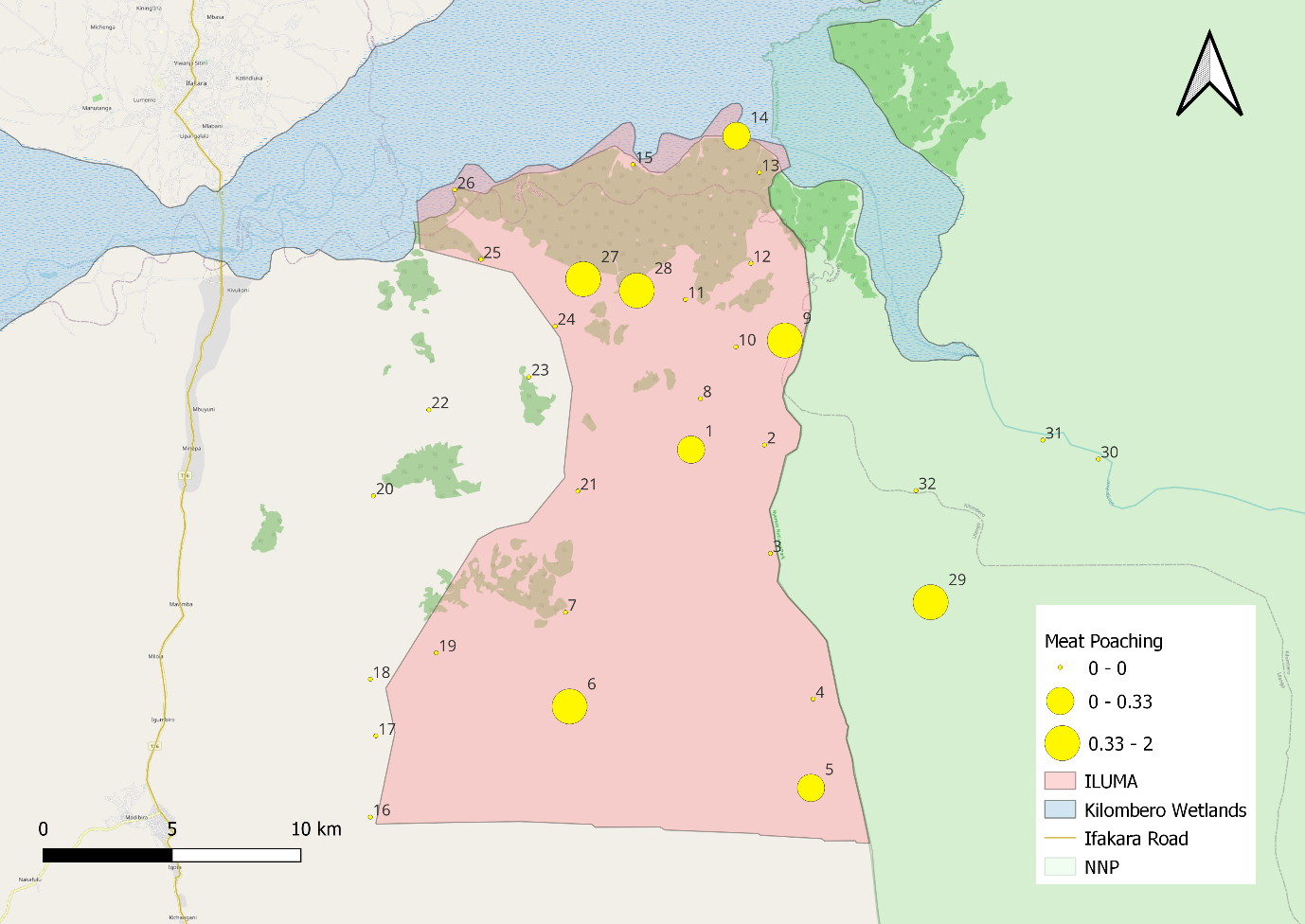

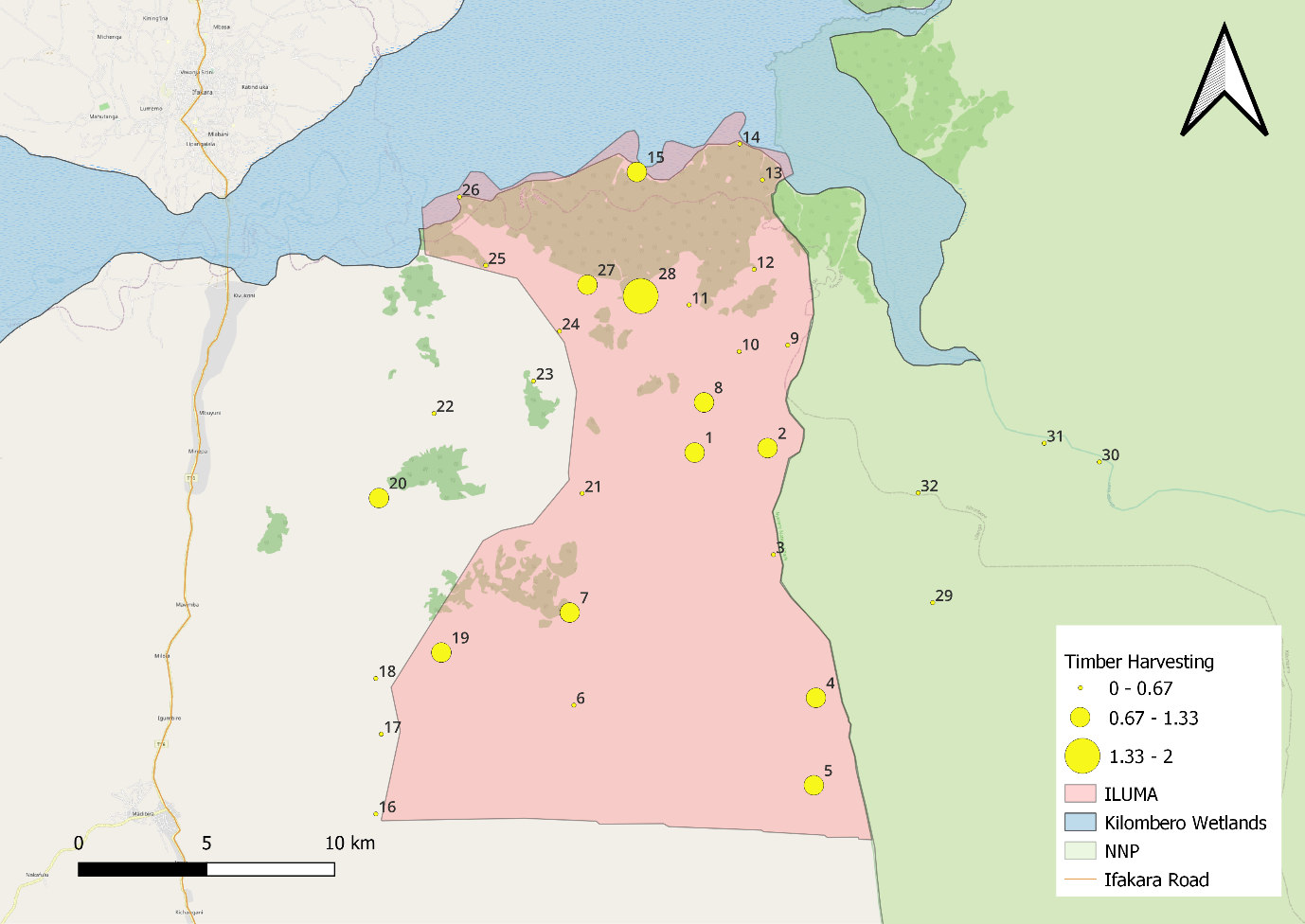

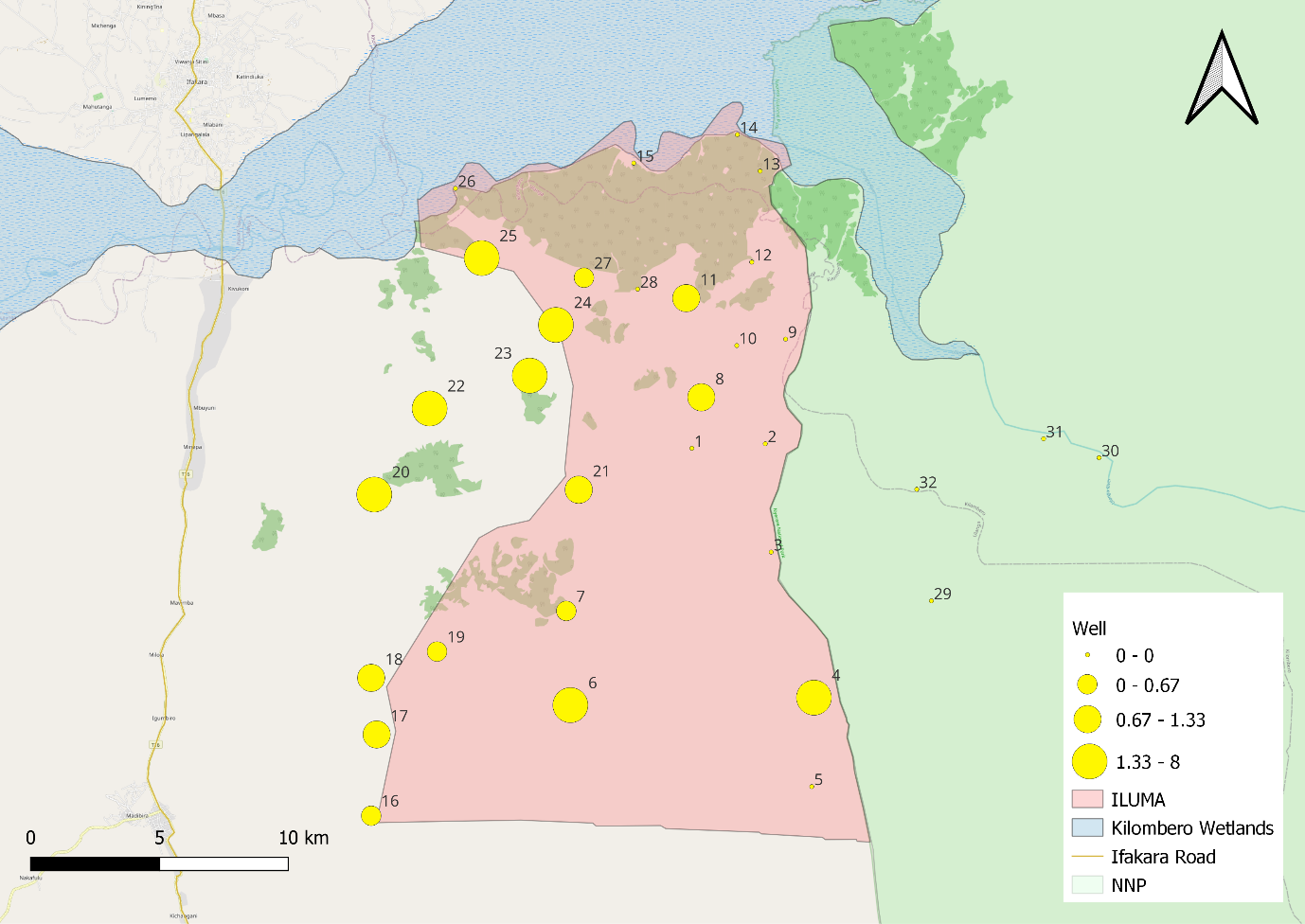

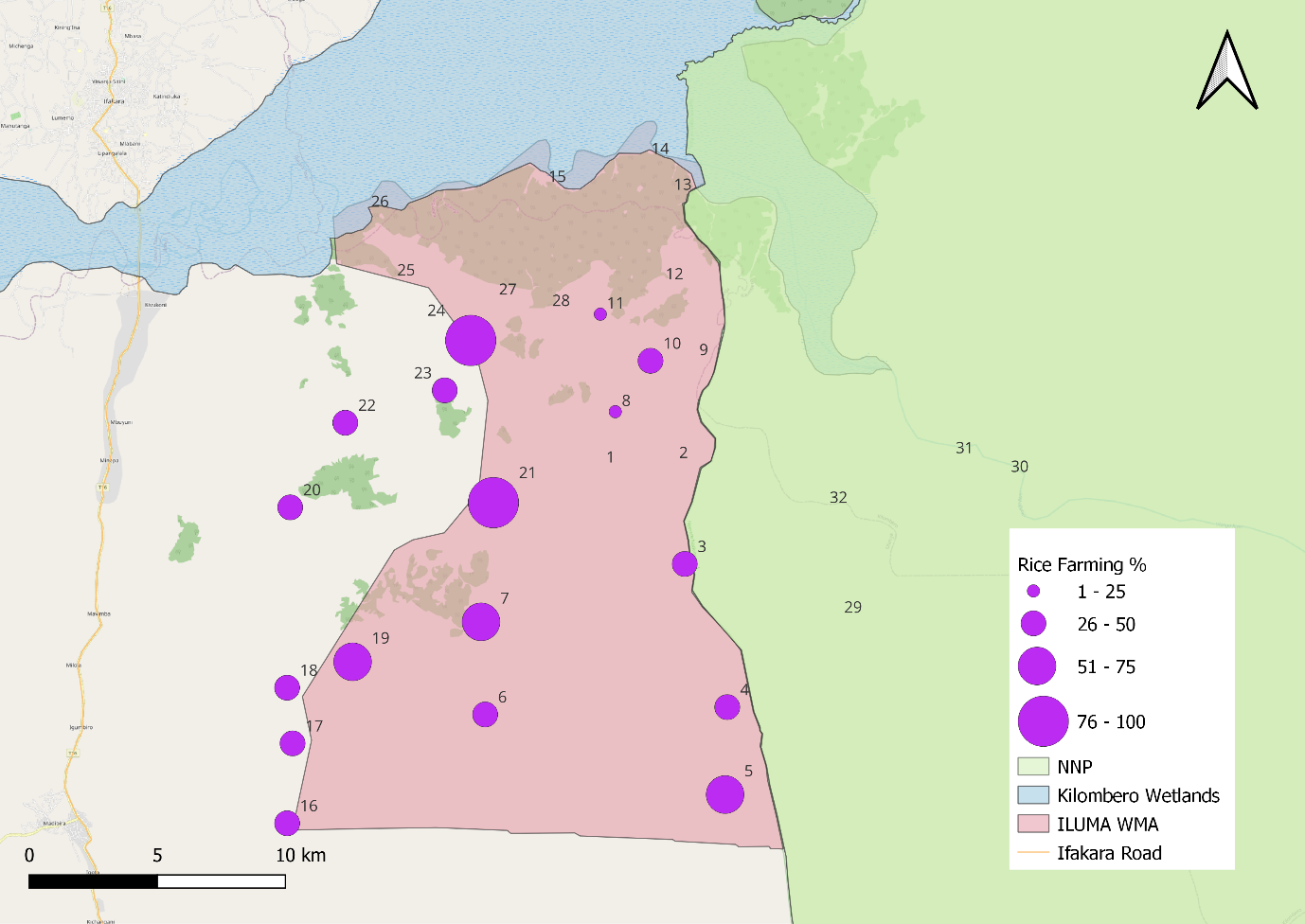

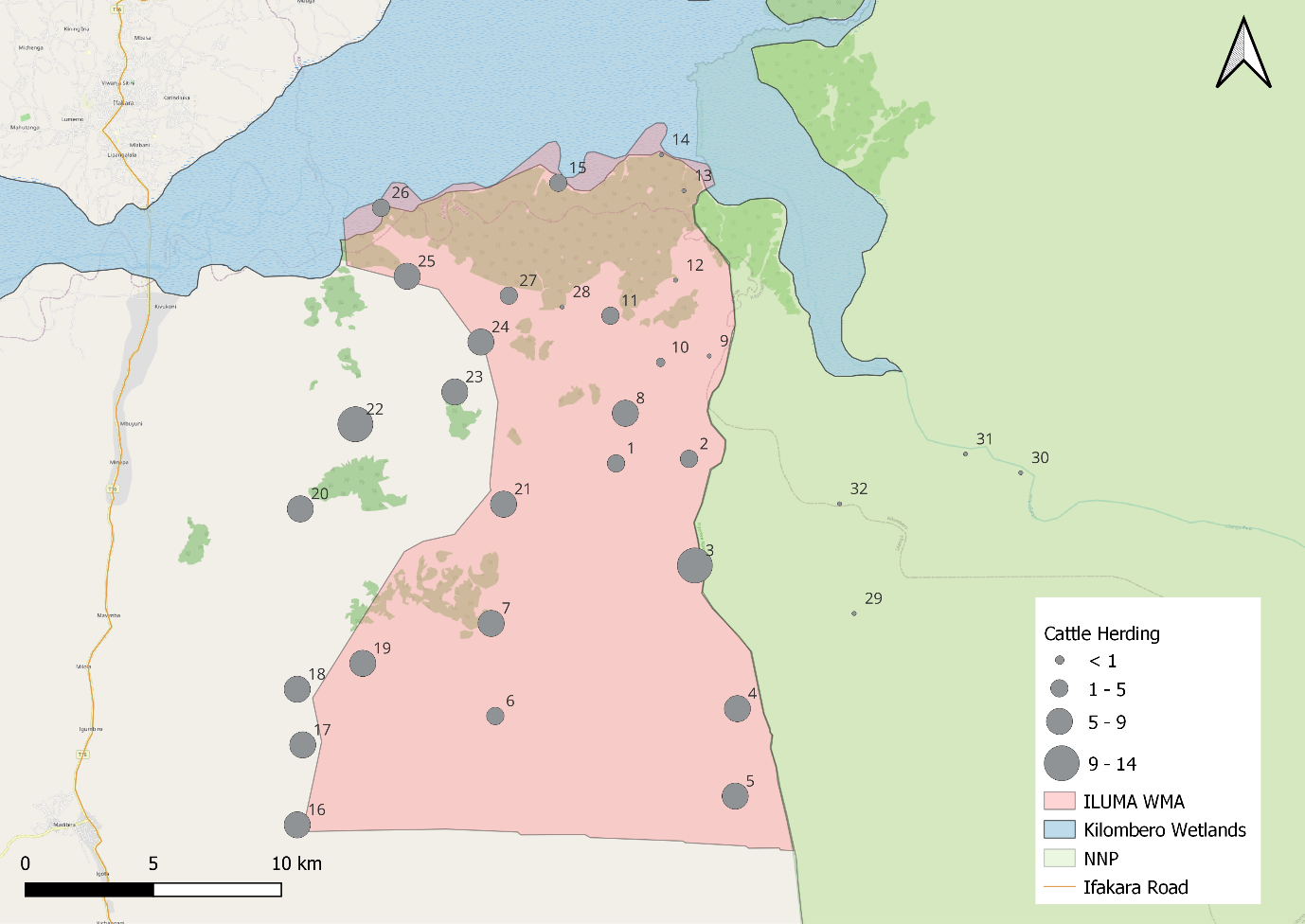

### Supplementary File 11

**Supp 11.** Maps illustrating significant human activities in final multivariate model.

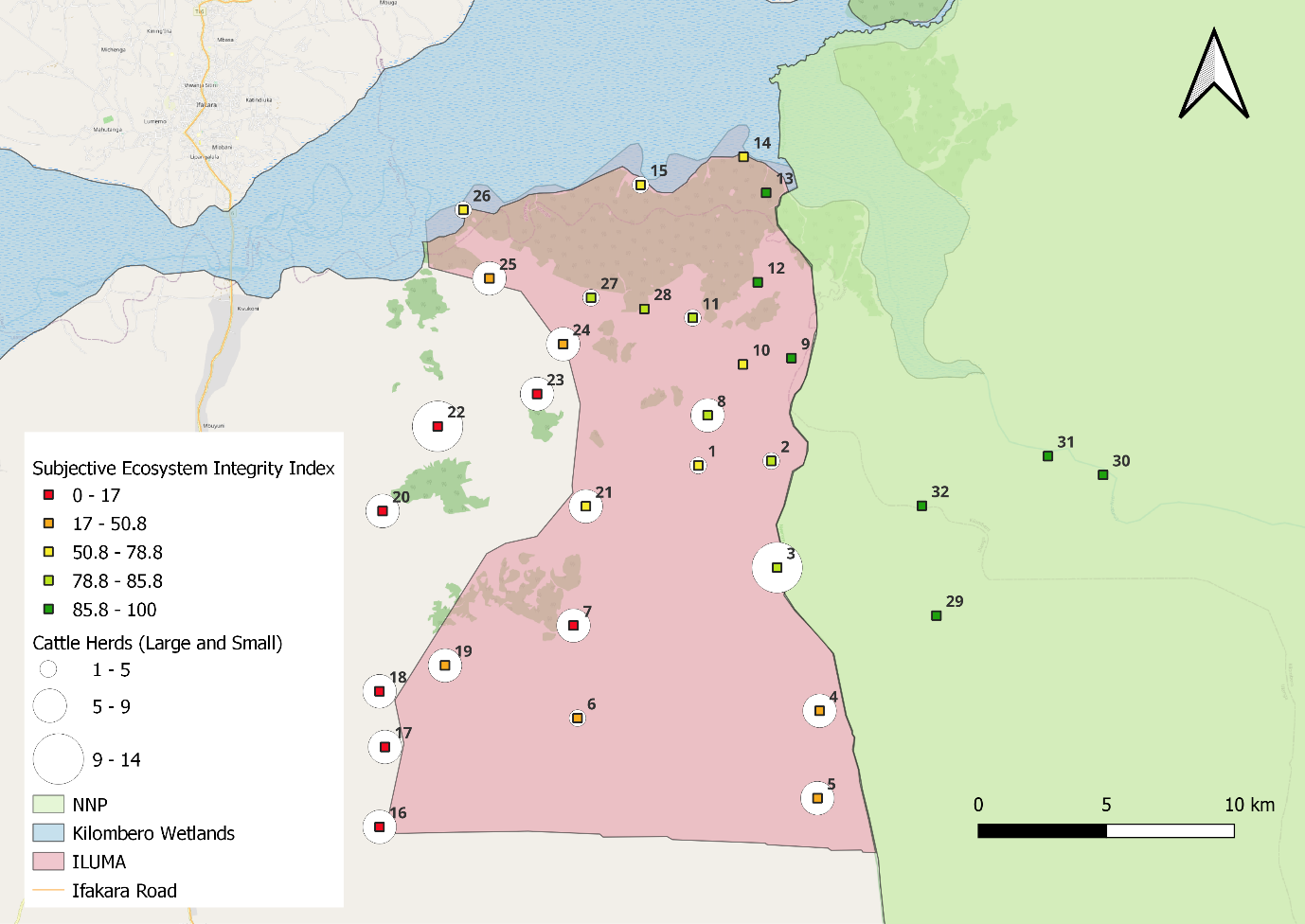

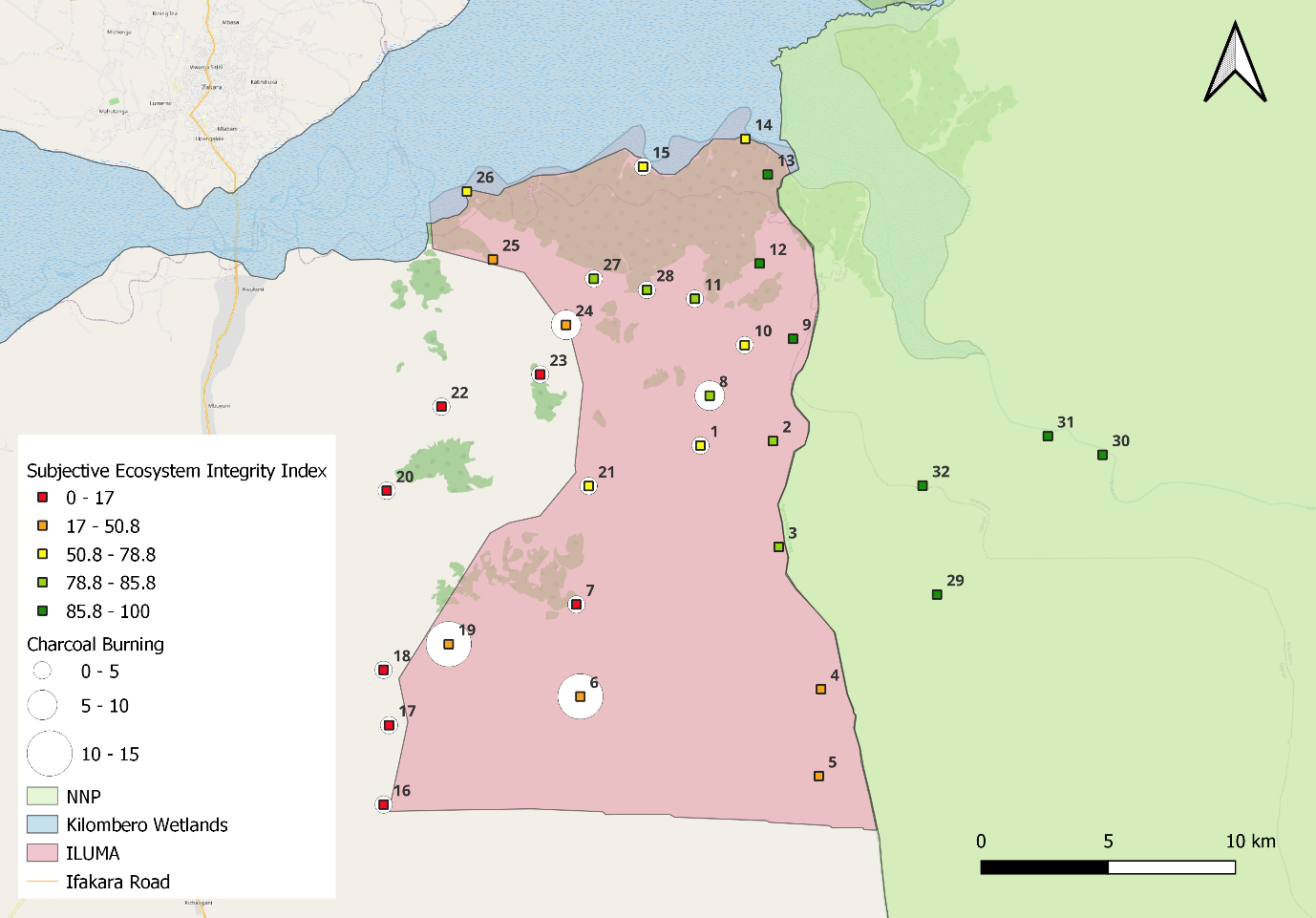

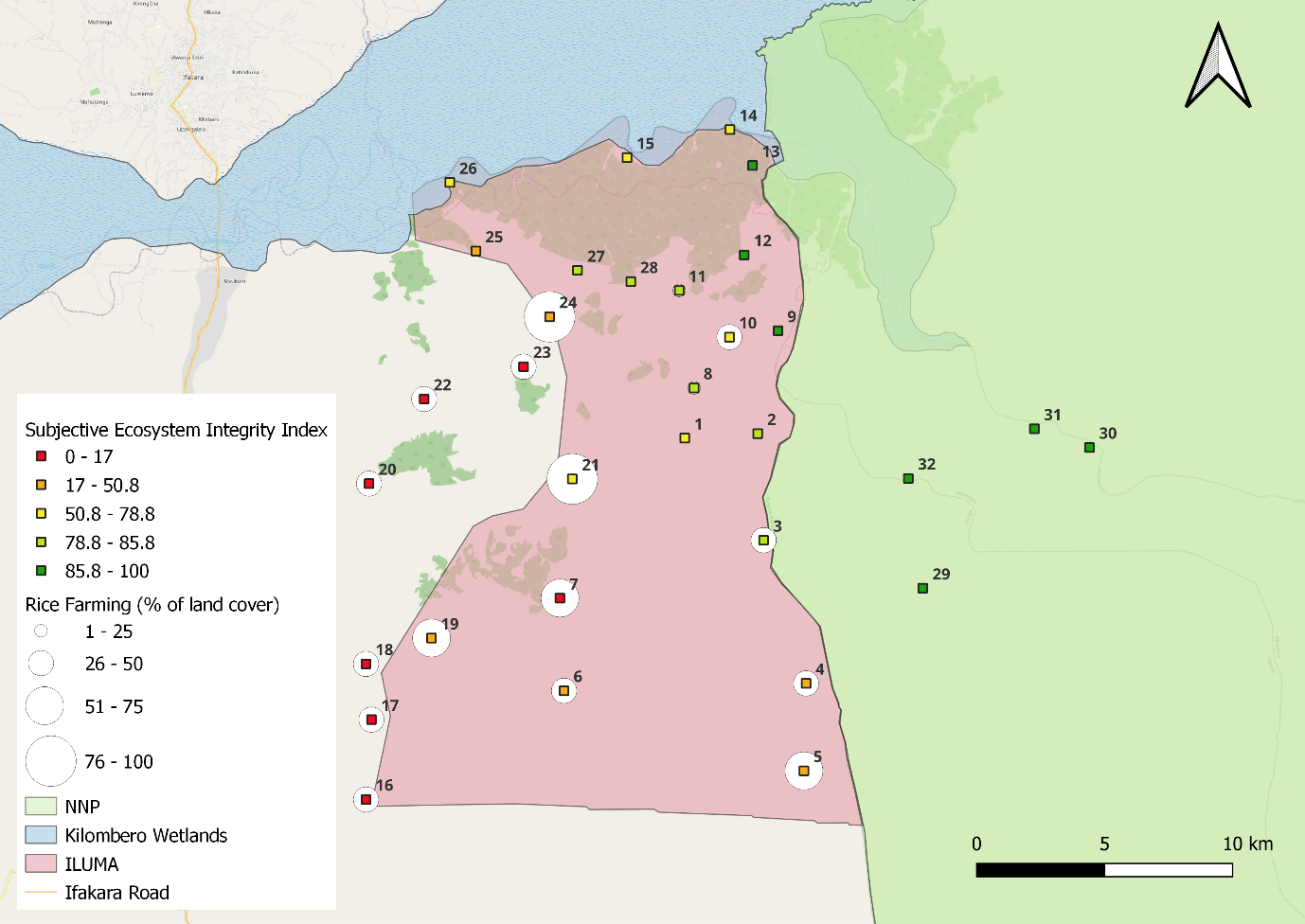

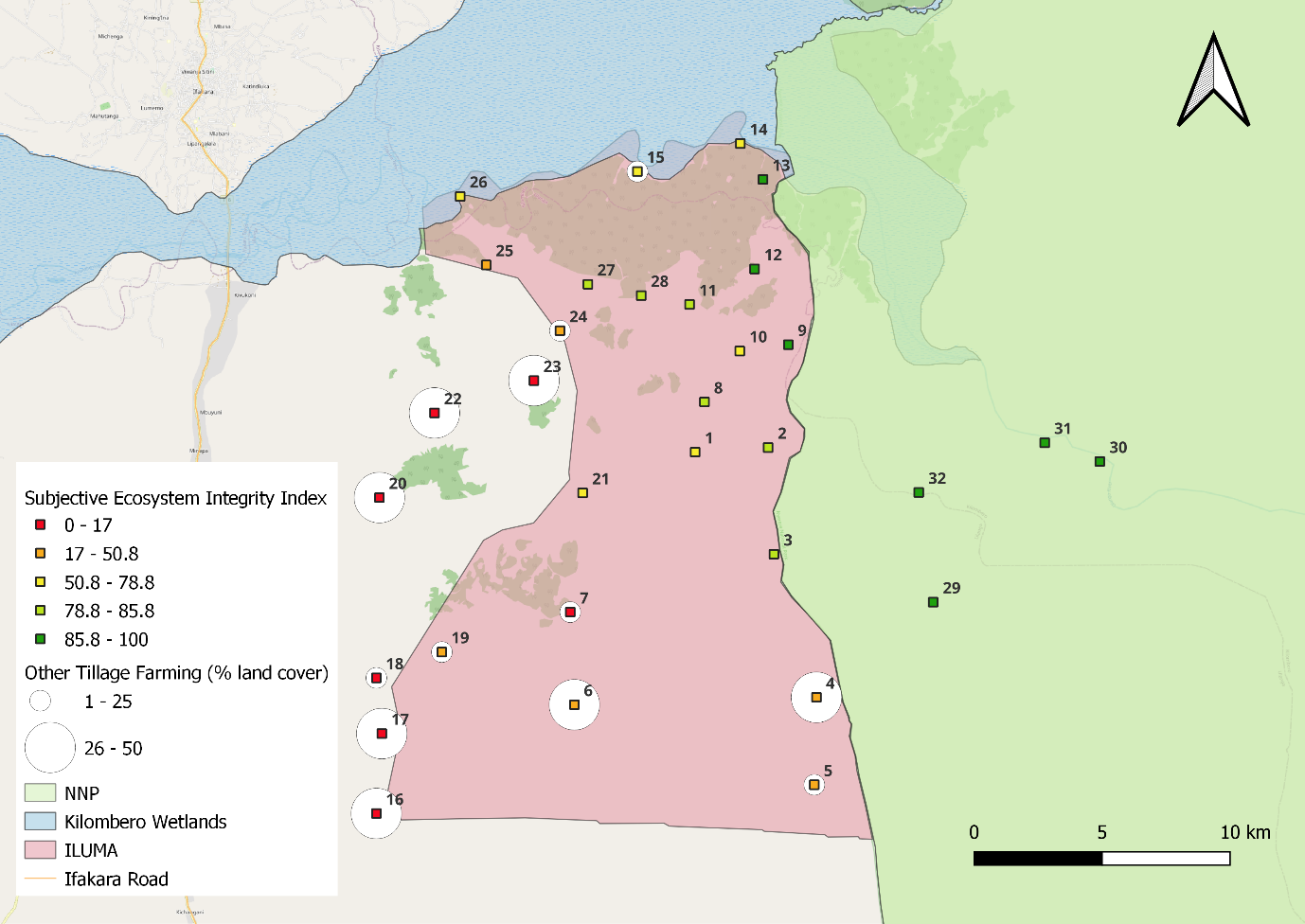

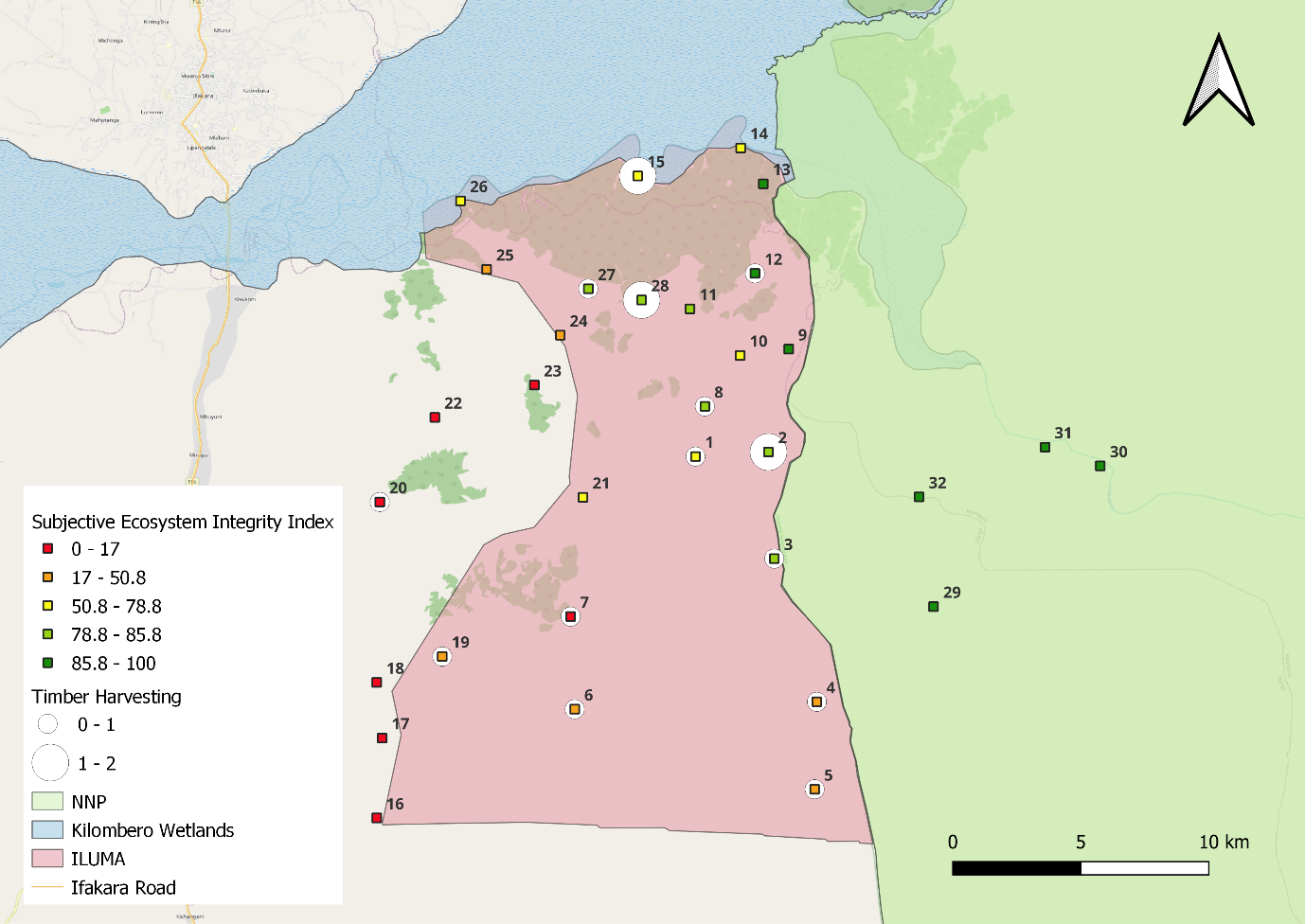
