## Supplementary File 10 for "An effective model for community-based conservation around authorized fishing settlements inside a devolved Wildlife Management Area in southern Tanzania" (from another version of this preprint: version 1, 2024-07-31)

**Table 10.1.** Results of univariate and multivariate generalized linear model (GLM) analyses of scaled human activity indicators as determinants of the estimated Simpson’s Index of Diversity Index (SID) for each camp surveyed^a^, with statistically significant associations highlighted in bold.

|  | Univariate | | | | Multivariate | | |
| --- | --- | --- | --- | --- | --- | --- | --- |
| Activity | Estimate **±** SEM | | t | P | Estimate **±** SEM | t | P |
| Livestock Herding | | **-0.2525 ± 0.0420** | **-6.01** | **<0.0001** | **-0.2183 ± 0.0513** | **-4.254** | **0.0002^b^** |
| Charcoal Burning | | -0.1134 **±** 0.0616 | -1.842 | 0.0754 | 0.0306 **±** 0.0363 | 0.842 | 0.4069^c^ |
| Timber Harvesting | | **-0.0889 ± 0.0433** | **-2.054** | **0.0488** | -0.0137 **±** 0.0254 | -0.540 | 0.5938^c^ |
| Fishing | | -0.0326 **±** 0.0467 | -0.697 | 0.4913 | **-0.0823 ± 0.0295** | **-2.788** | **0.0094^b^** |
| Hunting | | -0.0294 **±** 0.0447 | -0.659 | 0.5151 | 0.0044 **±** 0.0245 | 0.180 | 0.8586^c^ |
| Human Settlement | | **-0.1911 ± 0.0577** | **-3.314** | **0.0024** | -0.0455 **±** 0.0515 | -0.883 | 0.3850^c^ |
| Well | | **-0.2445 ± 0.0740** | **-3.306** | **0.0025** | 0.0522 **±** 0.0703 | 0.742 | 0.4642^c^ |
| Meat Poaching | | -0.0390 **±** 0.0437 | -0.893 | 0.379 | 0.0012 **±** 0.0252 | 0.047 | 0.9632^c^ |
| Human Presence | | -0.3654 **±** 0.0889 | -4.109 | **0.0003** | -0.1289 **±** 0.0742 | -1.739 | 0.0931^b^ |
| Rice Farming | | **-0.1788 ± 0.0436** | **-4.100** | **0.0003** | 0.0206 **±** 0.0414 | 0.498 | 0.6225^c^ |
| Other Tillage Farming | | **-0.2183 ± 0.0430** | **-5.078** | **<0.0001** | -0.0203 **±** 0.0512 | -0.396 | 0.6950^c^ |

SEM: Standard error of the mean.

^a^ The best fit models identified for this SID outcome assumed a Gaussian distribution and log link function for the dependent variable.

^b^ As estimated from the final, most parsimonious best-fit GLM, in which this was included on the basis of either being a significant (P≤0.05) determinant of SID or approached significance (P≤0.10) as a determinant of SID and significantly (P≤0.05) improved the goodness of fit based on the estimated Akaike Information Criterion (AIC).

^c^ As estimated from the last GLM in which this variable was still included before being removed on the basis that it was not a significant determinant of the SID (P>0.05) or approached significance (P≤0.10) but did not significantly improve the goodness of fit based on the estimated AIC (P>0.05).

**Table 10.2.** Results of univariate and multivariate linear regression analyses of scaled human activity indicators as determinants of the estimated Species Richness Index (SRI) for each camp surveyed^a^, with statistically significant associations highlighted in bold.

|  | Univariate | | | Multivariate | | |
| --- | --- | --- | --- | --- | --- | --- |
| Activity | Estimate **±** SEM | t | P | Estimate **±** SEM | t | P |
| Livestock Herding | **0.0186 ± 0.0056** | **3.348** | **0.0022** | **0.0186 ± 0.0056** | **3.348** | **0.0022^b^** |
| Charcoal Burning | **0.0273 ± 0.0115** | **2.381** | **0.0238** | 0.0038 **±** 0.0047 | 0.806 | 0.4271^c^ |
| Timber Harvesting | **0.0049 ± 0.0018** | **2.636** | **0.0132** | 0.0016 **±** 0.0010 | 1.508 | 0.1425^c^ |
| Fishing | 0.0000 **±** 0.0034 | -0.008 | 0.9940 | 0.0046 **±** 0.0028 | 1.649 | 0.1099^c^ |
| Hunting | **0.0030 ± 0.0010** | **2.923** | **0.0065** | 0.0010 **±** 0.0007 | 1.504 | 0.1434^c^ |
| Human Settlement | 0.0150 **±** 0.0080 | 1.871 | 0.0712 | 0.0017 **±** 0.0060 | 0.278 | 0.7830^c^ |
| Well | **0.0376 ± 0.0143** | **2.634** | **0.0132** | -0.0026 **±** 0.0065 | -0.404 | 0.6890^c^ |
| Meat Poaching | 0.0009 **±** 0.0026 | 0.336 | 0.7390 | -0.0015 **±** 0.0018 | -0.825 | 0.4160^c^ |
| Human Presence | 0.0667 **±** 0.0508 | 1.314 | 0.1989 | 0.0657 **±** 0.0633 | 1.037 | 0.3083^c^ |
| Rice Farming | **0.0117 ± 0.0040** | **2.940** | **0.0063** | 0.0031 **±** 0.0030 | 1.031 | 0.3111^c^ |
| Other Tillage Farming | **0.0132 ± 0.0051** | **2.587** | **0.0148** | 0.0033 **±** 0.0043 | 0.759 | 0.4543^c^ |

SD: Standard deviation of the coefficient point estimate.

P: P – Value.

^a^ The best fit models identified for this SRI outcome assumed a Gaussian distribution and inverse link function for the dependent variable, so positive values therefore indicate a negative association between the independent and dependent variable.

^b^ As estimated from the final, most parsimonious best-fit GLM, in which this was included on the basis of either being a significant (P≤0.05) determinant of SRI or approached significance (P≤0.10) as a determinant of SRI and significantly (P≤0.05) improved the goodness of fit based on the estimated Akaike Information Criterion (AIC).

^c^ As estimated from the last GLM in which this variable was still included before being removed on the basis that it was not a significant determinant of the SRI (P>0.05) or approached significance (P≤0.10) but did not significantly improve the goodness of fit based on the estimated AIC (P>0.05).

**Table 10.3.** Results of univariate and multivariate linear regression analyses of human activity indicators of scaled human activity indicators as determinants of thefor each camp surveyed^a^, with statistically significant associations highlighted in bold.

|  | Univariate | | | Multivariate | | |
| --- | --- | --- | --- | --- | --- | --- |
| Activity | Estimate **±** SEM | t | P | Estimate **±** SEM | t | P |
| Livestock Herding | **-0.5132 ± 0.1574** | **-3.275** | **0.0027** | **-0.4419 ± 0.1514** | **-2.918** | **0.0067^b^** |
| Charcoal Burning | -0.3775 **±** 0.1691 | -2.233 | 0.1691 | -0.1293 **±** 0.1722 | -0.751 | 0.4591 |
| Timber Harvesting | 0.0363 **±** 0.1825 | 0.199 | 0.844 | 0.1912 **±** 0.1543 | 1.239 | 0.2255 |
| Fishing | 0.2658 **±** 0.176 | 1.51 | 0.141 | 0.1357 **±** 0.1529 | 0.888 | 0.3822 |
| Hunting | -0.0618 **±** 0.1822 | -0.339 | 0.737 | -0.4365 **±** 0.1532 | -0.029 | 0.9775 |
| Human Settlement | **-0.3741 ± 0.1693** | **-2.21** | **0.0349** | -0.1306 **±** 0.1717 | -0.761 | 0.4532 |
| Well | -0.3463 **±** 0.1713 | -2.022 | 0.0522 | 0.1040 **±** 0.2311 | 0.450 | 0.6561 |
| Meat Poaching | **-0.4246 ± 0.1653** | **-2.568** | **0.0154** | **-0.3286 ± 0.1514** | **-2.170** | **0.0384^b^** |
| Human Presence | -0.2564 **±** 0.1765 | -1.453 | 0.157 | 0.2483 **±** 0.2215 | 1.121 | 0.2718 |
| Rice Farming | **-0.5066 ± 0.1574** | **-3.218** | **0.0031** | -0.2547 **±** 0.2063 | -1.235 | 0.2272 |
| Other Tillage Farming | **-0.465 ± 0.1616** | **-2.877** | **0.0073** | -0.1706 **±** 0.2223 | -0.767 | 0.4494 |

SD: Standard deviation of the coefficient point estimate.

P: P – Value.

^a^ The best fit models identified for this NACI outcome assumed a Gaussian distribution and identity link function for the dependent variable.

^b^ As estimated from the final, most parsimonious best-fit GLM, in which this was included on the basis of either being a significant (P≤0.05) determinant of SEII or approached significance (P≤0.10) as a determinant of NACI and significantly (P≤0.05) improved the goodness of fit based on the estimated Akaike Information Criterion (AIC).

^c^ As estimated from the last GLM in which this variable was still included before being removed on the basis that it was not a significant determinant of the NACI (P>0.05) or approached significance (P≤0.10) but did not significantly improve the goodness of fit based on the estimated AIC (P>0.05).
